# Tubular Mitochondrial Pyruvate Carrier Disruption Elicits Redox Adaptations that Protect from Acute Kidney Injury

**DOI:** 10.1101/2023.01.31.526492

**Authors:** Adam J. Rauckhorst, Gabriela Vasquez Martinez, Gabriel Mayoral Andrade, Hsiang Wen, Ji Young Kim, Aaron Simoni, Kranti A. Mapuskar, Prerna Rastogi, Emily J Steinbach, Michael L. McCormick, Bryan G. Allen, Navjot S. Pabla, Ashley R. Jackson, Mitchell C. Coleman, Douglas R. Spitz, Eric B. Taylor, Diana Zepeda-Orozco

**Affiliations:** Department of Molecular Physiology and Biophysics, University of Iowa, Iowa City, IA, USA; Fraternal Order of Eagles Diabetes Research Center (FOEDRC), University of Iowa, Iowa City, IA, USA; FOEDRC Metabolomics Core Research Facility, University of Iowa, Iowa City, IA, USA; Kidney and Urinary Tract Research Center, The Abigail Wexner Research Institute at Nationwide Children’s Hospital, Columbus OH, USA; Stead Family Department of Pediatrics, University of Iowa, Iowa City, IA, USA; Division of Pharmaceutics and Pharmacology, College of Pharmacy & Comprehensive Cancer Center, The Ohio State University, Columbus, OH, USA; Free Radical and Radiation Biology Program, Department of Radiation Oncology, University of Iowa, Iowa City, IA, USA; Holden Comprehensive Cancer Center, University of Iowa, Iowa City, IA, USA; Department of Pathology, University of Iowa, Iowa City, IA, USA; Department of Orthopedics and Rehabilitation, University of Iowa, Iowa City, IA, USA; Department of Pediatrics, The Ohio State University, Columbus, OH, USA; Pappajohn Biomedical Institute, University of Iowa, Iowa City, IA, USA

**Keywords:** acute kidney injury, mitochondrial metabolism, oxidative damage, metabolomics

## Abstract

Energy-intensive kidney reabsorption processes essential for normal whole-body function are maintained by tubular epithelial cell metabolism. Tubular metabolism changes markedly following acute kidney injury (AKI), but which changes are adaptive versus maladaptive remain poorly understood. In publicly available data sets, we noticed a consistent downregulation of the mitochondrial pyruvate carrier (MPC) after AKI, which we experimentally confirmed. To test the functional consequences of MPC downregulation, we generated novel tubular epithelial cell-specific *Mpc1* knockout (MPC TubKO) mice. ^13^C-glucose tracing, steady-state metabolomic profiling, and enzymatic activity assays revealed that MPC TubKO coordinately increased activities of the pentose phosphate pathway and the glutathione and thioredoxin oxidant defense systems. Following rhabdomyolysis-induced AKI, MPC TubKO decreased markers of kidney injury and oxidative damage and strikingly increased survival. Our findings suggest that decreased mitochondrial pyruvate uptake is a central adaptive response following AKI and raise the possibility of therapeutically modulating the MPC to attenuate AKI severity.

## INTRODUCTION

Acute kidney injury (AKI) is a major health problem (1, 2). In hospitalized patients, AKI is associated with increased length of stay, cost, and risk of mortality. Furthermore, survivors of AKI are at risk for greater injury from subsequent renal insults and for developing chronic kidney disease (3, 4). Tubular cell injury is an initiating event in the pathophysiological cascade of AKI. Although mechanisms of tubular injury are diverse, altered mitochondrial function and increased oxidative stress are common features. Tubular cells are rich in mitochondria that support ATP production to maintain their energy-intensive reabsorption processes. The high metabolic rate and mitochondrial content of tubular cells in-turn confers a high vulnerability to secondary oxidative injury following a primary insult (reviewed in (5-7)). Thus, understanding the relationship between tubular cell mitochondrial energetics and redox processes is critical to develop improved methods of attenuating AKI.

Mitochondrial pyruvate oxidation is a central feature of kidney metabolism, regulates redox balance, and is decreased during AKI (8-12). All tubular segments utilize pyruvate derived primarily from circulating lactate and secondarily from glycolysis for mitochondrial metabolism (13, 14). Conversion of lactate to pyruvate in route to mitochondrial oxidation generates cytosolic NADH, and oxidation of glycolytically produced pyruvate decreases glucose availability for NADPH production by the pentose phosphate pathway (PPP). Within mitochondria, pyruvate oxidation provides NADH and FADH2 that energize the electron transport chain (ETC) and drive oxidative phosphorylation for bulk ATP production. However, when mitochondria are dysfunctional, pyruvate oxidation can contribute to dysregulated ETC activity as a primary producer of cytotoxic reactive oxygen species (ROS). Under conditions of pathological ROS, channeling glucose into the PPP is cytoprotective by generating the NADPH required as cofactor for the glutathione and thioredoxin antioxidant systems. Thus, during AKI glucose, lactate, and pyruvate metabolism must be highly coordinated to prevent ROS-dependent cellular damage while maintaining adequate energy production. However, the mechanisms leading to the decreased mitochondrial pyruvate oxidation in and how they contribute to AKI are not well understood.

The mitochondrial pyruvate carrier (MPC) regulates the fate of glucose and lactate by transporting their common product pyruvate into mitochondria for TCA cycle oxidation. The MPC is a mitochondrial inner-membrane protein complex formed by two obligate subunits, MPC1 and MPC2 (15, 16). Studies of MPC disruption in diverse systems illustrate a conserved metabolic adaptive program where glucose and lactate oxidation decrease, glycolysis increases, and TCA cycle glutamine oxidation increases (17-24). In some cases, this contributes to disease, such as in many cancers where MPC disruption augments the Warburg effect (25, 26). Conversely, in others, MPC disruption is therapeutic, such as in adult skeletal muscle and liver, where it attenuates type 2 diabetes (27-30). Notably, in the liver, which like the kidney has profound capacity to oxidize glutamine, the increased glutamine oxidation caused by MPC disruption competes with glutathione synthesis and adaptively increases glutathione turnover through the transsulfuration pathway (31). Thus, the MPC occupies a nexus of metabolism, impacting energetics, substrate preference, and reactive oxygen species defense systems, and can be therapeutically disrupted in some tissues without collateral damage (27-30). However, the role of the MPC in normal kidney metabolism and in response to AKI, which is distinctively marked by decreased pyruvate oxidation and increased oxidative stress, is poorly defined.

Here, we address the role of the MPC in basic kidney tubule metabolism and AKI. Data from publicly available large-scale datasets corroborated by our own experiments showed that the MPC is downregulated after AKI, raising the question of how decreased MPC activity affects AKI severity. To investigate this, we generated novel kidney tubule epithelial cell-specific MPC knockout (MPC TubKO) mice and implemented a rhabdomyolysis model of AKI. MPC TubKO decreased glucose, lactate, and pyruvate oxidation in the TCA cycle, increased glucose flux through the distal PPP and NADPH levels, and increased glutathione turnover and reduction state. MPC TubKO strikingly increased survival from AKI, which was accompanied by decreased ROS-mediated tubular injury and increased glutathione and thioredoxin metabolism. Our findings demonstrate a central role of the kidney tubular cell MPC in metabolic regulation. They suggest that MPC deficiency coordinately protects from AKI by rewiring glucose metabolism to increase glycolysis and PPP activity and hormetically upregulating the glutathione and thioredoxin antioxidant systems. They provide a first example of modulating mitochondrial carbon fuel transport to protect from AKI, highlighting the potential therapeutic value of targeting mitochondrial transporters to protect from metabolic injury.

## RESULTS

### Mpc1 is expressed in proximal and distal tubular segments and is decreased in AKI

Mitochondrial pyruvate oxidation impacts cellular redox state through multiple pathways and is decreased in AKI (8-11). Because mitochondrial pyruvate uptake gates pyruvate oxidation, changes in tubular MPC activity could contribute to the decreased pyruvate oxidation and redox perturbations of AKI. To examine this, we first queried publicly available RNAseq datasets and observed tubular *Mpc1* mRNA abundance to be significantly downregulated following cisplatin-, ischemia reperfusion-, and rhabdomyolysis-induced AKI (**Supplemental Figure 1A, B**) (32, 33). To test the reproducibility of these results, we implemented similar mouse models of each. We found that, in addition to *Mpc1* mRNA, MPC1 protein levels were significantly decreased 72 hours after cisplatin-, 24 hours after ischemia-reperfusion-, and 24 hours after rhabdomyolysis-induced AKI (**Figure 1A, B**). Because rhabdomyolysis-induced AKI decreased MPC1 protein the most among the models tested, we deepened our analysis of rhabdomyolysis-injured tissue. Immunofluorescence analysis showed that tubular MPC1 protein was primarily decreased in the corticomedullary junction with a major decrease in the distal tubular segments (**Figure 1C**). We then generated mT/mG/Ggt1-Cre reporter mice that express membrane-localized GFP in renal tubular epithelial cells (RTECs), whereas all other cell-types express membrane-localized tdTomato (**Figure 1D**). RTECs showed significantly decreased *Mpc1* mRNA and MPC1 protein abundance following rhabdomyolysis-induced AKI, which remained unchanged in Non-RTECs (**Figure 1E, F**). VDAC protein levels were unaffected by injury, suggesting that the decreased RTEC MPC1 abundance occurred independent of changes in total mitochondrial content (**Figure 1F**).

**FIGURE 1.**
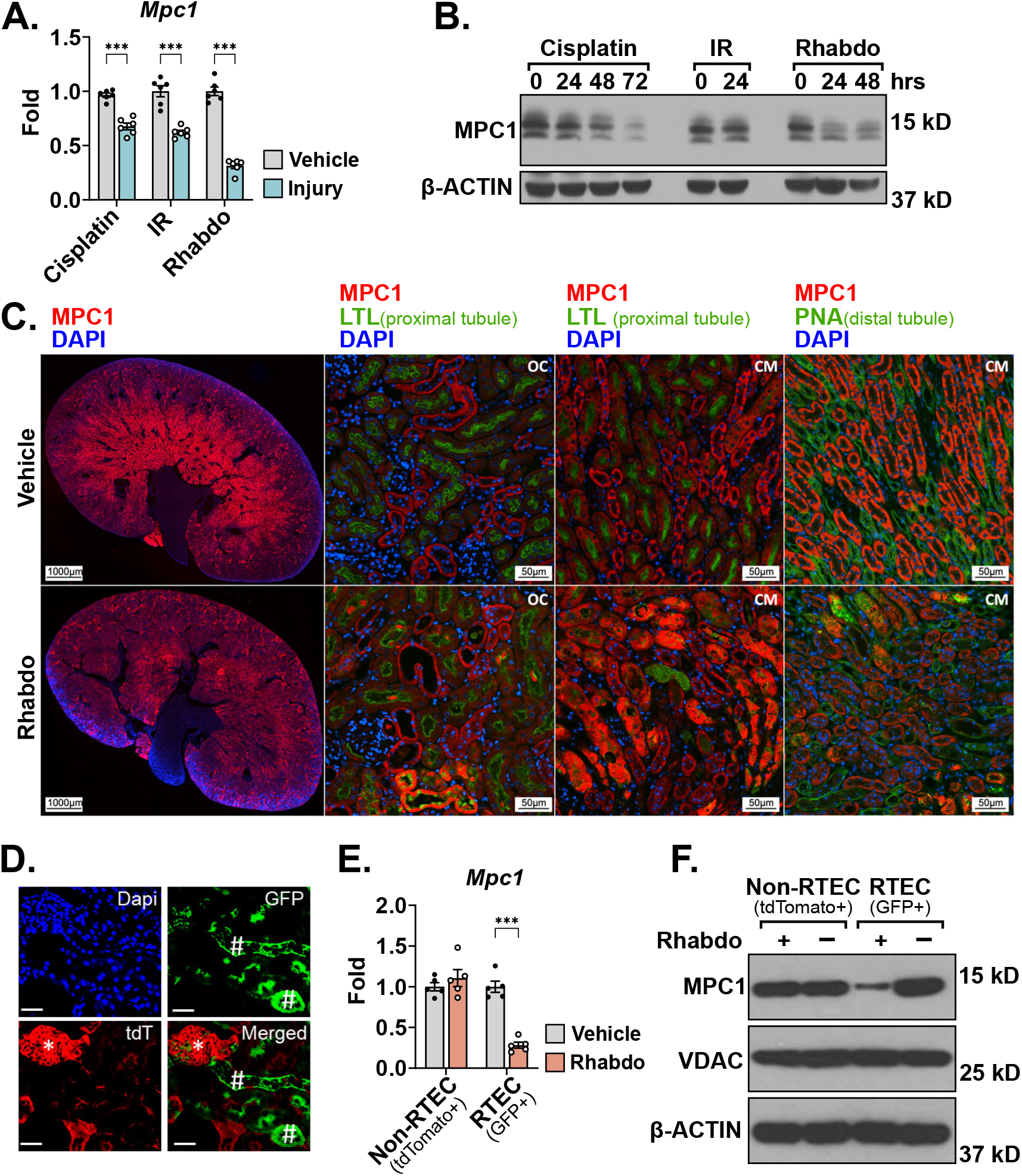
Mpc1 is downregulated in tubular epithelial cells during acute kidney injury. **(A)** Bar graph comparing kidney *Mpc1* mRNA levels after vehicle treatment or cisplatin-, ischemia reperfusion (IR)-, and rhabdomyolysis (Rhabdo)-induced AKIs. Samples were collected 72 hours after cisplatin injury and 24 hours after IR and rhabdomyolysis injuries. (n = 6/group; *** p < 0.001 by unpaired t test with Welch’s correction). **(B)** Representative Western blot of kidney MPC1 protein abundance after AKI. Samples were collected 72 hours after cisplatin injury and 24 hours after IR and rhabdomyolysis injuries. β-ACTIN was blotted as a loading control. **(C)** Representative immunostaining images of MPC1 (red), lotus tetragonolobus lectin (LTL, green, proximal tubule marker), or peanut agglutinin (PNA, green, distal tubule marker), and DAPI (blue) in whole kidney, outer cortex (OC) and cortico-medullary junction (CM) kidney sections 30 hours following vehicle treatment or rhabdomyolysis-induced AKI. (Images captured at 15x magnification; whole kidney scale bar = 1,000 μm; OC and CM scale bar = 50 μm). **(D)** Representative fluorescence image of kidney sections of mT/mG/Ggt1-Cre mice confirming GFP+ renal tubular epithelial cells (green, #, GFP) and tdTomato+ non-RTEC cells (red, *, tdT) stained with Dapi (blue). (Scale bar = 100 μm). **(E)** Bar graph comparing *Mpc1* mRNA levels in flow-sorted Non-RTEC (tdTomato+) and RTEC (GFP+) cells 24 hour after vehicle treatment or rhadbomyolysis-induced AKI. (n = 5/group, ***p < 0.001 by unpaired t test with Welch’s correction). **(F)** Representative Western blot of MPC1 and VDAC protein abundance in flow-sorted Non-RTEC (tdTomato+) and RTEC (GFP+) cells 24 hour after AKI. β-ACTIN was blotted as a loading control. Data are presented as means + SEM.

### Generation and characterization of MPC TubKO mice

To test the role of the MPC in the kidney tubule in vivo, we generated *Mpc1* pan-tubular epithelial cell knock out mice (MPC TubKO) by crossing *Mpc1*^f/f^ mice with *Pax8*-Cre mice (**Figure 2A**). MPC TubKO mice were viable at birth, and at 8 weeks of age they displayed normal body weight and levels of the renal function marker Cystatin C (**Figure 2B, C**). *Mpc1* mRNA was decreased by greater than 90% in MPC TubKO kidney tissue, which propagated to similar decreases in both MPC1 and MPC2 protein content (**Figure 2D-G**). We performed immunofluorescence staining across tubular segments to identify potential regions with residual MPC1 protein content (**Figure 2H**). Compared to WT controls, MPC1 was nearly absent in the proximal tubule cells of the renal cortex and corticomedullary junction and decreased but not eliminated in the corresponding distal tubules in MPC TubKO mice. Pax8 is reported to be expressed in the developing brain and liver (34, 35). To test for *Mpc1* deletion in brain and liver, we measured mRNA and protein abundance in these tissues from WT vs MPC TubKO mice. Compared to WT controls, *Mpc1* mRNA and MPC1 protein levels were similar in the brains and livers of MPC TubKO, consistent with tubular cell-delimited MPC knockout (**Supplemental Figure 2A-F**).

**FIGURE 2.**
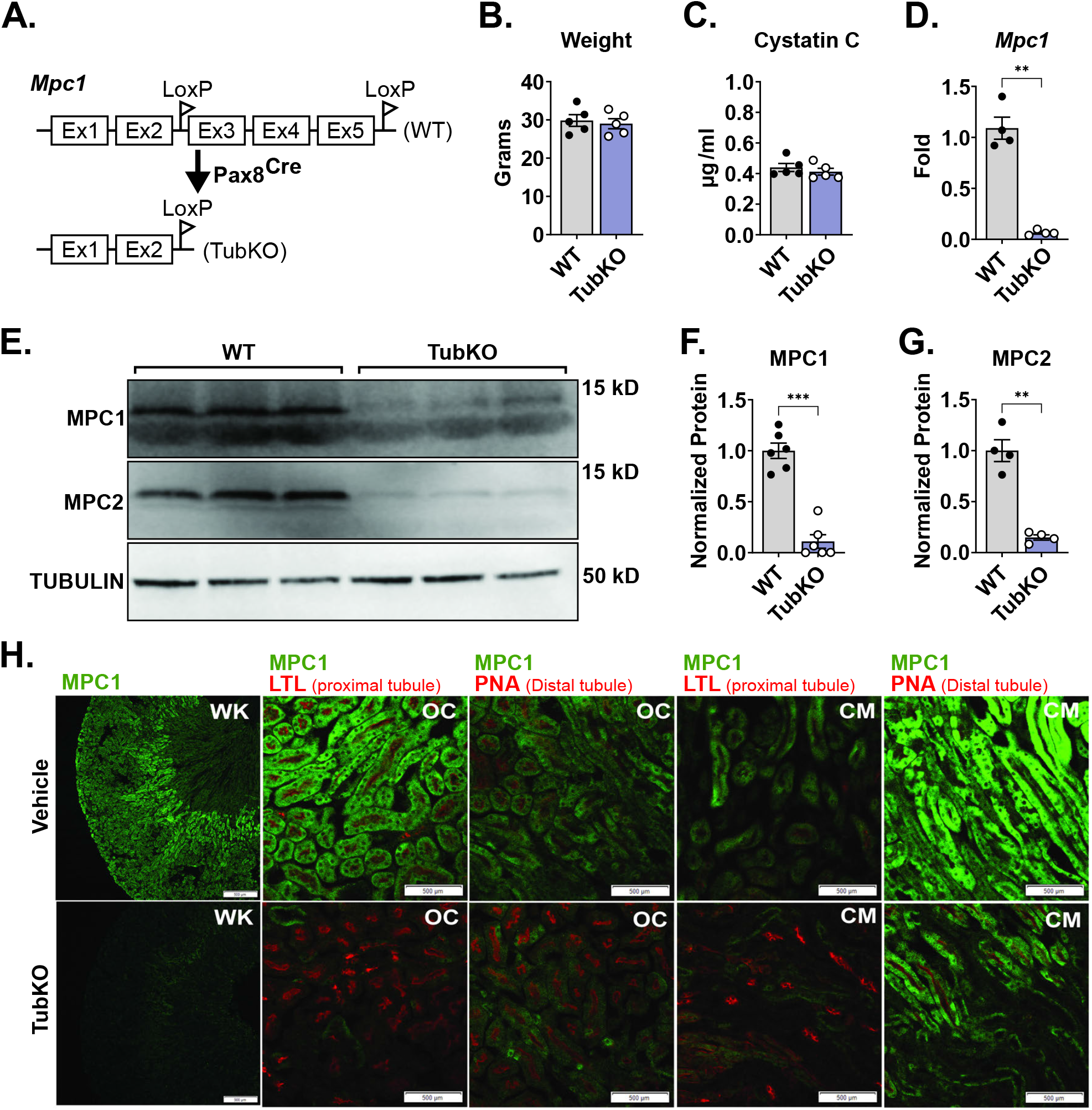
Generation and basic characterization of MPC TubKO mice. **(A)** Schematic illustrating the generation of tubular Mpc1 null allele, MPC TubKO mice (TubKO). **(B-C)** Bar graphs showing body weights (**B**) and serum cystatin C concentration (**C**) in WT and MPC TubKO mice. (n = 5/group, 8-week-old mice). **(D)** Bar graph comparing mouse kidney *Mpc1* mRNA levels in WT and MPC TubKO mice. (n = 4/group, 7 - 12-week-old mice, ** p < 0.01 by unpaired t test with Welch’s correction). **(E-G)** Representative Western blot of kidney MPC1 and MPC2 protein abundance (**E**) and quantification of normalized MPC1 (**F**) and MPC2 (**G**) levels in WT and MPC TubKO mice. Tubulin was blotted as loading control and used as the protein quantification normalizer. (n = 4 - 6/group, 7 - 12-week-old mice, *** p < 0.001 and ** p < 0.01 by by unpaired t test with Welch’s correction). **(H)** Representative immunostaining images of kidney MPC1 (green) and lotus tetragonolobus lectin (LTL, green, proximal tubule marker) or peanut agglutinin (PNA, green, distal tubule marker) in whole kidney (WK), outer-cortex (OC), and cortico-medullary junction (CM) in WT and MPC TubKO mice. (Images taken at 4x (WK) or 20x (OC and CM) magnification, scale bar = 500 μm). Data are presented as means + SEM.

### Tubular *Mpc1* deletion increases mitochondrial glutamate oxidation and perturbs ETC function

Next, we considered how tubular cell MPC loss bioenergetically affects MPC TubKO mice. In other systems, MPC loss adaptively increases glutamine metabolism to maintain mitochondrial metabolite levels and drive electron transport chain (ETC) conductance (18, 20). To test for this adaptation, we performed high resolution respirometry on mitochondria isolated from MPC TubKO and WT kidneys. Glutamate and malate were provided as the sole oxidative fuels. Basal and non-mitochondrial (rotenone-inhibited) oxygen consumption rates (OCR) between MPC TubKO and WT kidney mitochondria were not different. In contrast, FCCP-uncoupled respiration was increased in MPC TubKO kidney mitochondria, suggesting that tubular MPC loss adaptively increases capacity to oxidize glutamine (**Figure 3A**). We then assessed VDAC protein levels and citrate synthase activity in whole kidney extracts as markers of mitochondrial content and found no differences between MPC TubKO mice and WT controls (**Figure 3B-D, Supplemental Figure 3A**). Similarly, no apparent differences were detected in markers of ETC protein abundance (**Figure 3B**). Given that each ETC complex comprises multiple subunits and that individual ETC protein levels do not denote enzymatic activities, we biochemically evaluated the activities of Complex I, II, and III. Complex I and complex II activities were similar between MPC TubKO and WT controls (**Figure 3E, F**). However, MPC TubKO complex III activity was significantly decreased suggesting that MPC loss may impair the ETC, which could increase ROS production (**Figure 3G**) (36). Together, these data suggest tubular MPC disruption directly affects mitochondrial function without altering cellular mitochondrial content.

**FIGURE 3.**
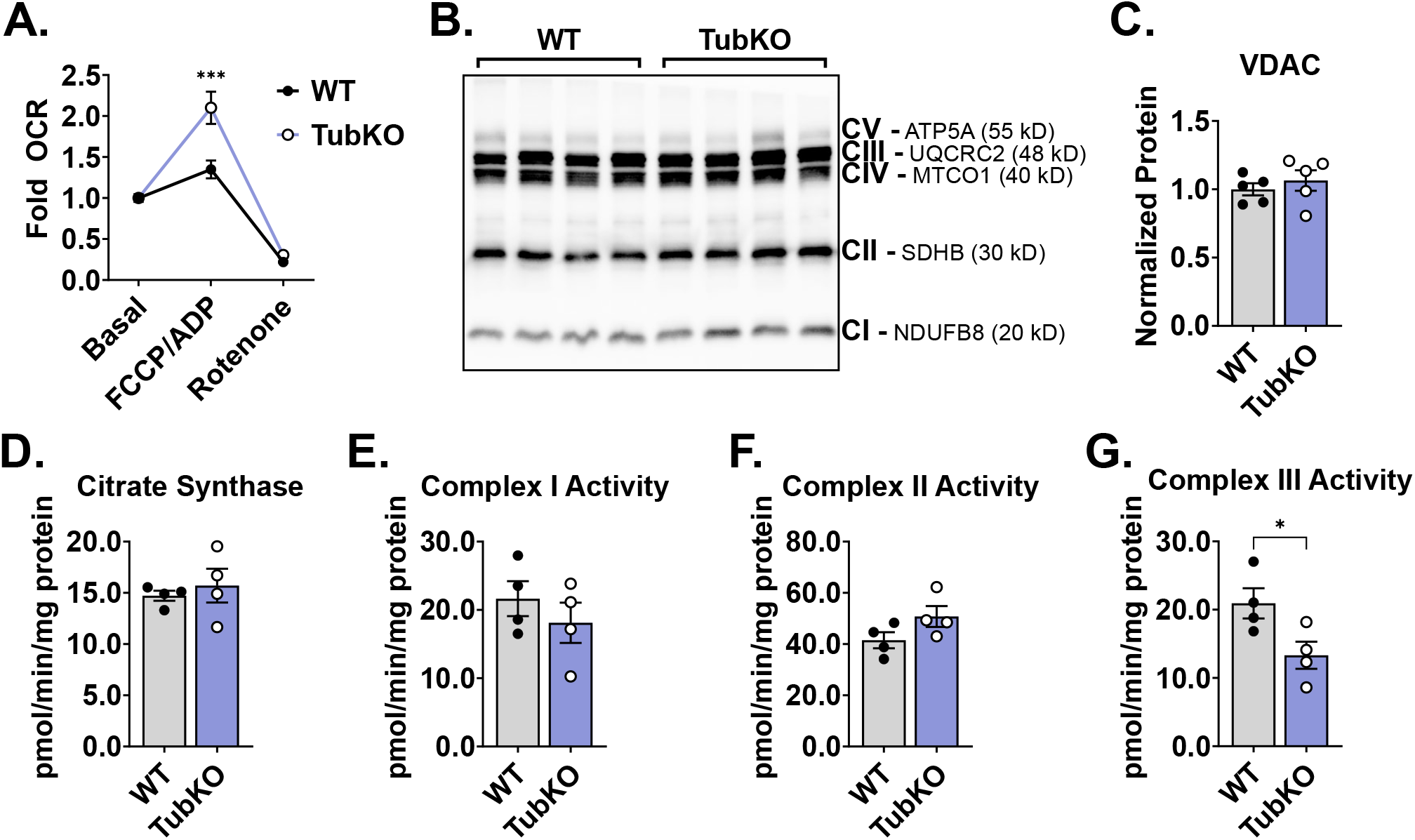
MPC TubKO mice have altered mitochondrial function. **(A)** Line graph showing the relative glutamate-fueled oxygen consumption rate (OCR) under basal (no addition), maximal FCCP/ADP-stimulated, and rotenone-inhibition conditions in kidney mitochondria isolated from WT and MPC TubKO mice. (n = 6/group, 8 - 12-week-old mice, *** p < 0.001 by two-way ANOVA with Tukey’s multiple comparison test). **(B)** Representative Western blot of kidney ETC marker Complex I (CI), NDUFB8; Complex II (CII), SDHB; Complex III (CIII), UQCRC2; Complex IV (CIV), MTCO1; and Complex V (CV), ATP5A protein abundances in WT and MPC TubKO mice. (n = 4/group, 6 - 8-week-old mice). **(C)** Bar graph comparing the quantified VDAC protein level in WT and MPC TubKO mice. (n = 5/group, 6 - 8-week-old mice) **(D)** Bar graph showing the whole-kidney citrate synthase enzymatic activity in WT and MPC TubKO mice. (n = 4/group, 6-week-old mice). (**E-G**) Bar graphs comparing the whole-kidney enzymatic activities of Complex I (**E**), Complex II (**F**), and Complex III (**G**) in WT and MPC TubKO mice. (n = 4/group, 6-week-old mice, * p < 0.05 by unpaired t test with Welch’s correction). Data are presented as means + SEM.

### Tubular MPC disruption leads to upregulation of oxidant defense systems

To identify metabolic changes evoked by tubular MPC disruption, we compared the steady state metabolomic profiles of freeze-clamped WT and MPC TubKO kidney tissue. The relative abundances of 49 metabolites were significantly different in MPC TubKO kidneys (**Figure S4A, Supplemental Table 3**). While pyruvate was not significantly increased, lactate and alanine, which are produced from pyruvate by single metabolic reactions and were previously identified to be increased in models of MPC disruption (28, 29, 37), were significantly increased in MPC TubKO kidneys (**Figure 4A**). TCA cycle metabolites were broadly decreased except for α-ketoglutarate, which is consistent with decreased pyruvate and increased glutamine oxidation (**Figure 4B**). Notably, MPC TubKO kidneys had decreased levels of the glutathione (GSH) synthesis substrates glycine, cysteine, and glutamate (**Figure 4C**) and increased levels of 2-hydroxybutyrate, a marker of glutathione turnover (31, 38, 39) (**Figure 4D**). Glutathione is the most abundant cellular antioxidant, a key determinant of redox signaling, a modulator of cell fate and function (reviewed in (40)), and an integral component for hydroperoxide and electrophile detoxification (**Figure 4E**). Because our metabolomic profiling data showed changes in glutathione metabolism, we measured total and oxidized glutathione (GSSG) levels using an enzyme-coupled reaction in whole kidney lysates. Total glutathione levels (GSH + GSSG) were unchanged in MPC TubKO mice (**Figure 4F**); however, the amount of GSSG in the total glutathione pool (GSH + GSSG) was increased in the MPC TubKO mice, consistent with increased glutathione synthesis and turnover (**Figure 4G**).

**FIGURE 4.**
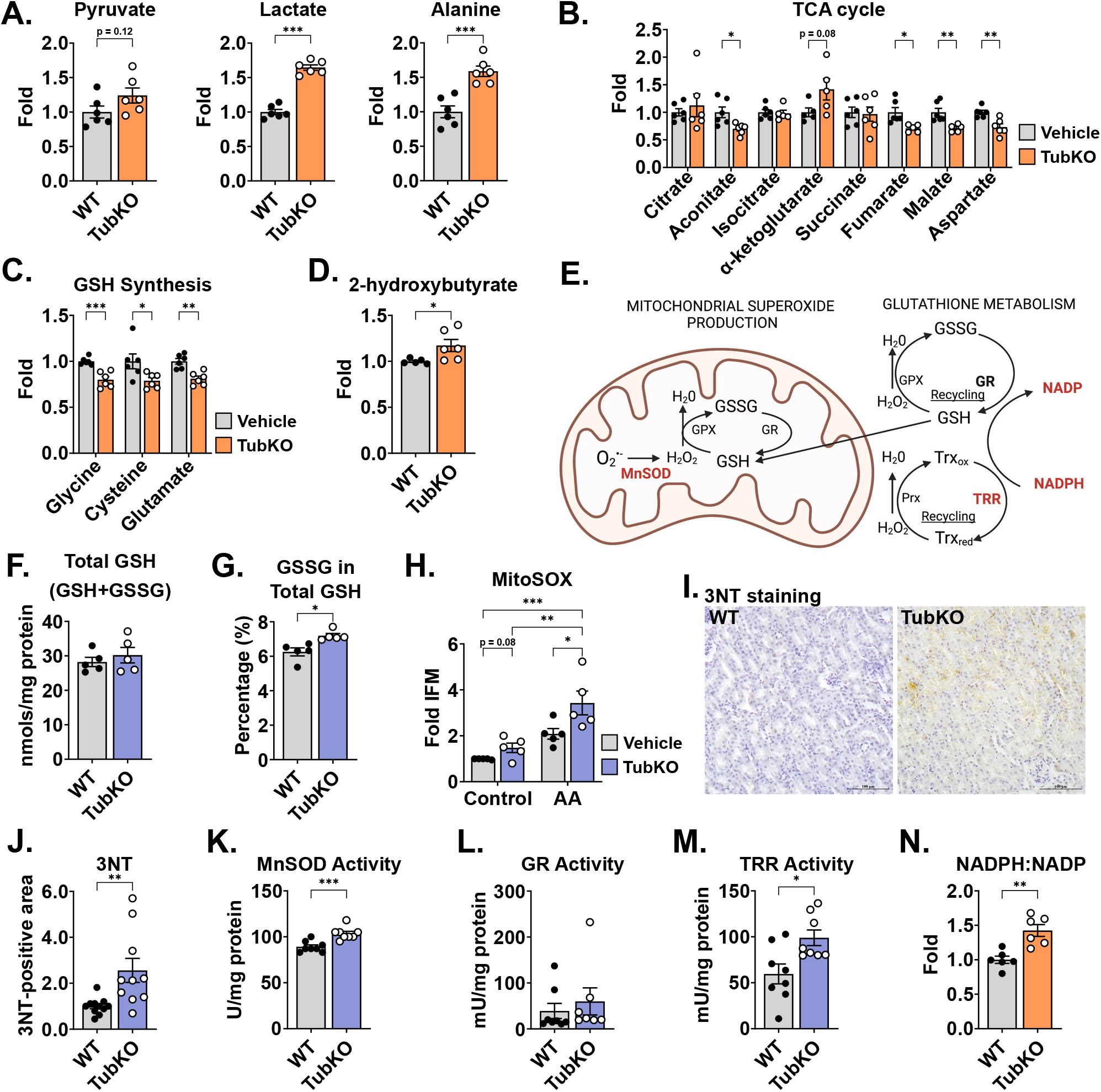
MPC TubKO mice have mitochondrial redox adaptation. (**A-D**) Bar graphs showing kidney metabolite levels in WT and MPC TubKO mice. Pyruvate, lactate, and alanine (**A**), TCA cycle metabolites (**B**), the GSH synthesis substrates glycine, cysteine, and glutamate (**C**), and 2-hydroxybutyrate, a marker of GSH turnover (**D**). (n = 6/group, 8 - 12-week-old mice, * p < 0.05, ** p < 0.01, and *** p < 0.001 by unpaired t test with Welch’s correction). **E**. Schematic illustrating mitochondrial antioxidant defense system including MnSOD, manganese superoxide dismutase; GSH, glutathione; GSSG, oxidized glutathione; Gpx, glutathione peroxidase; GR, glutathione reductase; Prx, peroxiredoxin; Trx, thioredoxin reductase; Trx_red_, reduced thioredoxin; and Trx_ox_, and oxidized thioredoxin. (**F-G**). Bar graphs comparing kidney total GSH (GSH + GSSG) (**F**) and the % of GSSG of total GSH (GSH + GSSG) (**G**) in WT and MPC TubKO mice. (n = 5/group, 12 - 14-week-old mice, * p < 0.05 by unpaired t test with Welch’s correction). **(H)** Bar graph showing MitoSOX oxidation in the presence and absence of antimycin A (AA) of isolated WT and MPC TubKO tubular epithelial cell. (n = 5/group, 12-week-old mice, * p < 0.05, ** p < 0.01, *** p < 0.001 by two-way ANOVA with Tukey’s multiple comparison test). **(I)** Representative immunohistochemistry images of kidney 3NT staining in WT and MPC TubKO mice. (Images taken at 40x magnification, scale bar = 100 μm). **(J)** Bar graph comparing kidney 3NT quantification in WT and MPC TubKO mice. (n = 8 - 11, 12 - 14-week-old mice, ** p < 0.01 by unpaired t test with Welch’s correction). (**K-M**) Bar graphs showing kidney enzyme activities of MnSOD (**K**), GR (**L**), and TRR (**M**) in WT and MPC TubKO mice. (n = 7 - 8/group, 12 - 14-week-old mice, * p < 0.05 and *** p < 0.001 by unpaired t test with Welch’s correction). (**N**) Bar graph comparing the kidney NADPH:NADP ratio in WT and MPC TubKO mice. (n = 6/group, 8 - 12-week-old mice, ** p < 0.01 by unpaired t test with Welch’s correction). Data are presented as means + SEM.

Our finding that MPC TubKO mice have decreased complex III activity and an altered GSH reduction state (**Figure 3G**) led us to speculate that loss of the tubular MPC could increase ROS levels and alter cellular redox homeostasis. To test this, we isolated primary tubular cells from MPC TubKO and WT mice and measured mitochondrial ROS production. Superoxide-dependent MitoSOX oxidation in MPC TubKO tubular cells trended towards increased under basal conditions and was enhanced more significantly by the complex III inhibitor antimycin A (**Figure 4H**). Next, we measured the kidney levels of 3-nitrotyrosine (3NT), which is formed from peroxynitrite (ONOO-: the product of O_2_•-+ •NO) reacted with tyrosine residues and is a representative marker of oxidative protein modification. MPC TubKO mice had increased 3NT levels (**Figure 4I, J**). These data demonstrate that that tubular MPC loss increases reactive oxygen species production and oxidative damage.

We expanded our studies to the effects of tubular MPC loss to mitochondrial redox response and cellular oxidant defense systems. As a first step of oxidant defense, the aberrant ETC-generated superoxide undergoes dismutation to hydrogen peroxide by manganese superoxide dismutase (MnSOD; aka SOD2) (**Figure 4E**). MPC TubKO mice had increased MnSOD activity compared to WT controls (**Figure 4K**). Next, we looked downstream to the GSH regenerating activity of glutathione reductase (GR), which reduces hydrogen peroxide to water thereby detoxifying ROS. GR activity was slightly, but not significantly, increased in MPC TubKO mice (**Figure 4L**, p = 0.15). Thioredoxin (Trx) provides a second thiol redox couple (Trx_ox_-Trx_red_) sustained by thioredoxin reductase (TRR), which maintains cellular hydrogen peroxide levels in parallel to GR (**Figure 4E**). Thioredoxin reductase activity was significantly increased in MPC TubKO mice (**Figure 4M**). Glutathione reductase and thioredoxin reductase catalysis require NADPH oxidation to NADP (**Figure 4E**). Consistent with increased support for glutathione reductase and thioredoxin reductase activity, we found increased NADPH and, inversely, decreased NADP with MPC TubKO (**Figure S4B**) resulting in an increased NADPH/NADP ratio in MPC TubKO mice (**Figure 4N**). This suggests that tubular MPC disruption leads to coordinated increases in NADPH redox cycling and activation of the glutathione and thioredoxin oxidant defense systems.

### MPC TubKO mice have increased pentose phosphate pathway activity

The pentose phosphate pathway (PPP) is the major source of NADPH regeneration in many systems. Thus, we considered whether increased PPP activity could contribute to the increased NADPH/NADP ratio and help sustain the increased activities of the glutathione and thioredoxin antioxidant systems in MPC TubKO mice. To test this, we examined ^13^C enrichments into PPP metabolites from U-^13^C-glucose as an indirect measure of NADPH production (**Figure 5A**). There were no apparent differences in ^13^C enrichments into glucose 6-phosphate and 6-phosphogluconate between MPC TubKO and WT mice (**Figure 5B, C, Supplemental Table 4)**. However, m+5 and total ^13^C enrichments into Ribo/Ribulose 5-phosphate were increased in MPC TubKO (**Figure 5D**). These data suggested that tubular MPC loss adaptively increased distal PPP activity to preserve an increased NADPH/NADP ratio supporting the glutathione reductase and thioredoxin reductase reactions and producing ribose sugars needed for DNA repair.

**FIGURE 5.**
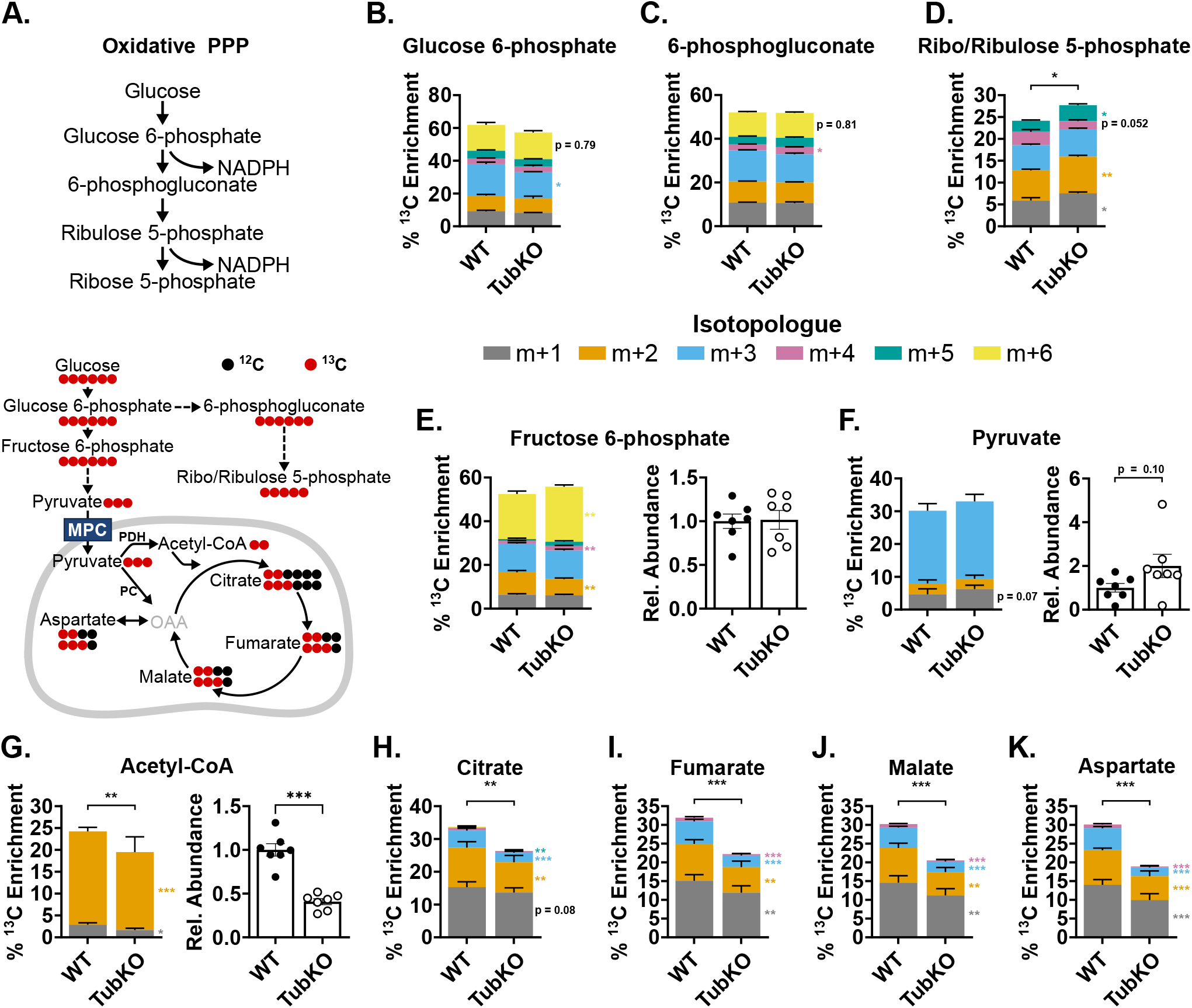
^13^C-glucose tracing shows increased distal PPP activity in MPC TubKO mice. (**A**) Schematics illustrating pentose phosphate pathway (PPP) (top) and ^13^C-enrichment patterns into glycolysis, the PPP, and the TCA cycle from ^13^C-glucose (bottom). MPC, mitochondrial pyruvate carrier; PDH, pyruvate dehydrogenase; PC, pyruvate carboxylase; OAA, oxaloacetate. (**B-D**) Stacked bar graphs showing kidney ^13^C-isotopologue enrichments into PPP metabolites 30 minutes after ^13^C-glucose bolus injection in WT and MPC TubKO mice. Glucose 6-phosphate (**B**), 6-phosphogluconate (**C**), and ribo/ribulose 5-phosphate (**D**). (n = 7/group, 7 - 8-week-old mice, * p < 0.05 and ** p < 0.01 by unpaired t test). (**E-G**) Stacked bar graphs showing kidney ^13^C-isotopologue enrichments and bar graphs comparing relative abundances of metabolites 30 minutes after ^13^C-glucose bolus injection in WT and MPC TubKO mice. Fructose 6-phosphate (**E**), pyruvate (**F**), and acetyl-CoA (**G**). (n = 7/group, 7 - 8-week-old mice, * p < 0.05 and ** p < 0.01 by unpaired t test). (**H-K**) Stacked bar graphs showing kidney ^13^C-isotopologue enrichments into TCA cycle metabolites 30 minutes after ^13^C-glucose bolus injection in WT and MPC TubKO mice. Citrate (**H**), Fumarate (**I**), Malate (**J**), and Aspartate as a surrogate measure of oxaloacetate (**K**). (n = 7/group, 7 - 8-week-old mice, * p < 0.05 and ** p < 0.01 by unpaired t test). Data are presented as means + SEM.

To evaluate the impact of MPC TubKO in the overall renal glucose handling, we examined ^13^C enrichments into fructose 6-phosphate, a metabolite of proximal glycolysis downstream of PPP shunting; pyruvate, the terminal glycolytic metabolite; and TCA cycle intermediates 30 minutes after a bolus injection of U-^13^C-glucose. M+6 fructose 6-phosphate ^13^C enrichments were significantly increased although total abundance was unchanged in MPC TubKO (**Figure 5E**), and ^13^C enrichment into pyruvate was similar in MPC TubKO mice. Total pyruvate abundance trended towards being increased in MPC TubKO mice, similar to what was observed when ^13^C lactate and pyruvate were administered (**Figure 5F, Supplemental Figure 5A, Supplemental Table 5**). ^13^C enrichment into and abundance of acetyl-CoA was decreased in MPC TubKO mice (**Figure 5G**) indicating that tubular MPC disruption limits glucose-driven TCA cycle metabolism. Indeed, decreased, but not eliminated, pyruvate oxidation in MPC TubKO mice resulted in decreased ^13^C enrichments into citrate, fumarate, and malate as we observed when ^13^C-lactate and -pyruvate were administered (**Figure 5H-J, Supplemental Figure 5B, C**). We also observed significantly decreased ^13^C enrichment into aspartate in MPC TubKO mice, which is a marker of oxaloacetate due to their rapid equilibration across the glutamate-oxaloacetate transamination reaction (**Figure 5K**). Residual TCA cycle intermediate ^13^C enrichments in MPC TubKO likely resulted from alanine-bypass activity as demonstrated in other MPC loss models (27, 28, 41). While the kidney has a role in systemic glucose homeostasis via gluconeogenesis and glucose reabsorption, tubular MPC deletion did not impact systemic blood glucose or lactate levels, following an 18 hour fast or in response to bolus injection of lactate/pyruvate (**Supplemental Figure 5D-I**). These data indicate that tubular MPC loss limits TCA cycle activity, increases glycolysis, and increases glucose flux through the distal oxidative PPP without affecting systemic glucose homeostasis.

### MPC TubKO mice are protected from ROS mediated damage

Because tubular MPC loss upregulated the glutathione and thioredoxin antioxidant systems and increased PPP activity, we next sought to understand how MPC TubKO impacts the PPP enzymatic activity response to AKI. We again utilized rhabdomyolysis-induced AKI, where ROS-dependent injury causes pan-tubular damage (**Figure 6A**) (42). First, we examined the activities of the NADPH-producing PPP enzymes glucose 6-phosphate dehydrogenase (G6PD) and 6-phosphogluconate dehydrogenase (6PGDH) (**Figure 6B**). Twenty-four hours after rhabdomyolysis injury, compared to WT controls, MPC TubKO kidney G6PD activity was significantly upregulated (**Figure 6C**) and 6PGDH trended to be upregulated (p = 0.07, **Figure 6D**). To test if NADPH utilizing oxidant defense systems were coordinately upregulated, we examined the activities of the glutathione reductase and thioredoxin reductase. Indeed, following injury, MPC TubKO mice more robustly upregulated glutathione reductase and thioredoxin reductase activities (**Figure 6E, F**). We then examined tubular protein glutathionylation as a stable marker of cellular oxidative stress (43, 44), which was blunted in MPC TubKO mice following AKI (**Figure 6G, H**). This shows that mice lacking tubular MPC activity more resiliently maintain redox homeostasis following AKI. Together, these data provide an enzyme-level mechanistic basis for how tubular MPC loss-dependent PPP upregulation supports increased glutathione and thioredoxin antioxidant systems during rhabdomyolysis-induced AKI.

**FIGURE 6.**
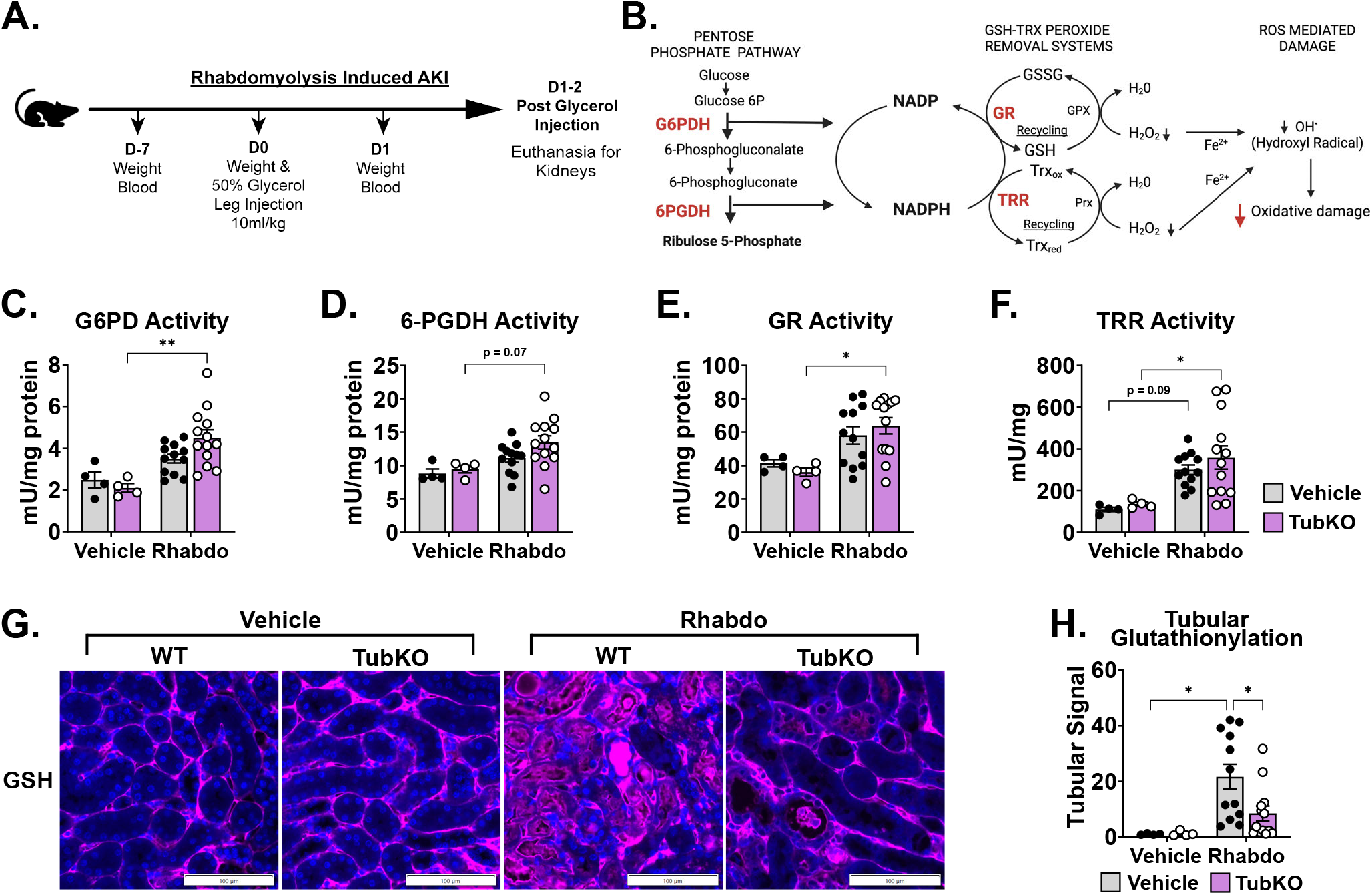
Downregulation of tubular Mpc1 is an early adaptive response to protect from oxidative damage. (**A-B**) Schematics illustrating the time course of the rhabdomyolysis-induced AKI model (**A**) and the interconnectedness of the pentose pathway and cellular antioxidant defense systems (**B**). (**C-F**) Bar graphs showing kidney enzyme activities following vehicle treatment or rhabdomyolysis (Rhabdo)-induced AKI. Glucose-6-phosphate dehydrogenase (**C**, G6PD), 6- phosphogluconate dehydrogenase (**D**, 6PGDH), glutathione reductase (**E**, GR), and thioredoxin reductase (**F**, TRR). (n = 4/group for vehicle treatment, n = 12 - 13/group for Rhabdo, 8 - 12- week-old mice, * p < 0.05 and ** p < 0.01 by two-way ANOVA with Tukey’s multiple comparison test). **(F)** Representative immunostaining images of kidney protein-glutathionylation (pink) and Dapi (blue) following vehicle treatment or rhabdomyolysis-induced AKI in WT and MPC TubKO mice. (Scale bar = 100 μm, n = 4/group for vehicle treatment, n = 12 - 13/group for Rhabdo, 8 - 12- week-old mice). **(AG** Bar graph showing quantified kidney protein-glutathionylation following vehicle treatment or rhabdomyolysis-induced AKI in WT and MPC TubKO mice. (n = 4/group for vehicle treatment, n = 12 - 13/group for Rhabdo, 8 - 12-week-old mice, * p < 0.05 by two-way ANOVA with Tukey’s multiple comparison test). Data presented as means + SEM

### MPC TubKO mice are protected from rhabdomyolysis induced kidney injury

Finally, we extended our investigation to test the effect of MPC TubKO in clinical-translation outcomes following rhabdomyolysis-induced AKI. MPC TubKO markedly increased overall survival through 48 hours of injury (100% vs 63.6%, p = 0.05, n = 11 WT and 9 MPC TubKO) (**Figure 7A**). Furthermore, the renal function markers cystatin C and blood urea nitrogen (BUN), which were similar before injury, were lower in MPC TubKO mice 24 and 48 hours after AKI (**Figure 7B, C**). MPC TubKO mice similarly showed decreased transcript levels of the kidney tubular injury markers *Ngal* and *Kim1* 24 hours after AKI (**Figure 7D,E**), and the histologically assessed tubular injury score 24 hours after AKI was decreased (**Figure 7F**). As basic measure of cellular stress, we observed decreased tubular apoptosis assessed by tunel staining in MPC TubKO mice 24 hours after injury (**Figure 7G**). These results demonstrate that tubular MPC loss improves multiple markers of kidney injury and protects against rhabdomyolysis-induced tubular injury and AKI mortality.

**FIGURE 7.**
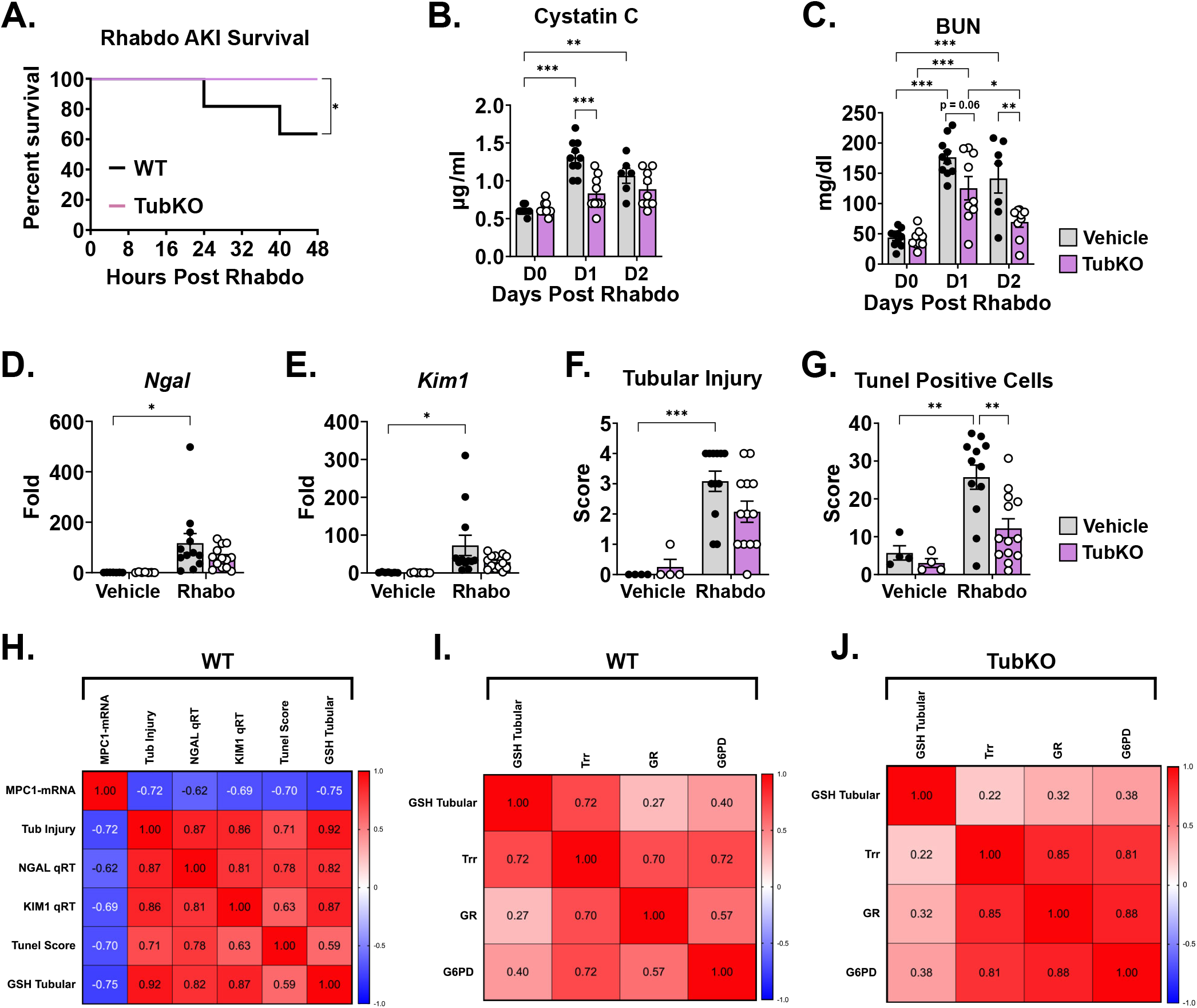
Tubular MPC1 genetic deletion protects from rhabdomyolysis induced kidney injury. (**A**) Line graph showing the survival curve of WT and MPC TubKO mice following rhabdomyolysis (Rhabdo)-induced AKI. (n = 10 - 11/group, 8 - 12-week-old mice, * p < 0.05 by Mantel-Cox log-rank test). (**B**-**C**) Bar graphs showing serum cystatin C (**C**), and blood urea nitrogen (**D**, BUN) levels prior to (D0) and on day 1 (D1, 24-hours) and day 2 (D2, 48 hours) after vehicle treatment or rhabdomyolysis-induced AKI in WT and MPC TubKO mice. (n = 10 - 11/group, 8 - 12-week-old mice, * p< 0.05, ** p < 0.01, *** p < 0.001 by two-way ANOVA followed by Turkey’s multiple comparison tests). (**E-F**) Bar graphs showing kidney *Ngal* (**D**) and *Kim1* (**E**) mRNA levels one day (24 hours) after vehicle treatment or rhabdomyolysis-induced AKI in WT and MPC TubKO mice. (n = 4/group for vehicle treatment, n = 12 - 13/group for Rhabdo, 8 - 12-week-old mice, * p < 0.05 by two-way ANOVA with Tukey’s multiple comparison test). (**G-H**) Bar graphs showing quantification of histologically assessed tubular injury score (**F**) and tunel positive tubular cells (**G**) one day (24 hours) after vehicle treatment or rhabdomyolysis-induced AKI in WT and MPC TubKO mice. (n = 4/group for vehicle treatment, n = 12 - 13/group for Rhabdo, 8 - 12-week-old mice, ** p < 0.01 and *** p < 0.001 by two-way ANOVA with Tukey’s multiple comparison test). (**H-J**) Heatmaps showing Spearman correlation between variables analyzed following vehicle treatment or rhabdomyolysis-induced AKI. Correlation calculated in WT mice comparing *Mpc1* mRNA levels, tubular injury, *Ngal* and *Kim1* mRNA levels, tunel score, and tubular GSH with AKI (**H**). Spearman correlation performed in WT (**I**) and MPC TubKO (**J**) mice comparing GSH and antioxidant defense system markers following rhabdomyolysis-induced AKI. Data are presented as means + SEM.

To statistically summarize how MPC TubKO modulates the glutathione and thioredoxin antioxidants systems during AKI, we correlated all rhabdomyolysis-induced AKI injury data. In WT mice, the degree of *Mpc1* mRNA decrease inversely correlated with the degree of tubular GSH (r = -0.75, p = 0.001), tubular injury (r = -0.72, p = 0.002), tunel score (r = -0.70, p = 0.004), and tubular injury biomarkers *Ngal* (r = -0.62, p = 0.0125) and *Kim1* (r = -0.69, p = 0.004) mRNA abundance (**Figure 7H**). Thioredoxin reductase activity positively correlated with tubular GSH in WT mice (r = 0.72, p = 0.002) which was lost in MPC TubKO mice (r = 0.22, p = 0.4) (**Figure 7I, J**). Glucose 6-phosphate dehydrogenase activity and thioredoxin reductase and glutathione reductase more strongly correlated in MPC TubKO mice (r = 0.81 and 0.88, p < 0.001) compared to WT controls (r = 0.72 and 0.57, p = 0.002 and 0.024 respectively) showing that MPC TubKO mice have an increased capacity to maintain tubular redox state and increased PPP activity following AKI.

## DISCUSSION

Kidney metabolic changes following AKI including increased glycolysis and decreased mitochondrial pyruvate oxidation are well described (8-11). However, the mechanisms regulating mitochondrial pyruvate oxidation after AKI and how pyruvate oxidation capacity affects AKI severity are not well defined. We found that MPC1 mRNA and protein levels are decreased in renal tubular epithelial cells (RTECs) following cisplatin-, ischemia reperfusion-, and rhabdomyolysis-induced AKI. Given this observation, we aimed to test the contribution of the tubular MPC to basic kidney metabolism and AKI severity. Our overall findings demonstrate that the MPC regulates tubular mitochondrial pyruvate oxidation in vivo and that MPC disruption induces upregulation of antioxidant systems and protects from AKI.

Our work demonstrates that the MPC plays a central role in tubular metabolism that is also dispensable for normal kidney function. Given the enormous energetic demands required for kidney function, and the decreased pyruvate oxidation observed after AKI, a key question is whether decreased pyruvate oxidation impairs kidney function. Tubular epithelial cells rely on mitochondrial activity to perform a variety of functions including ATP production, solute reabsorption, gluconeogenesis, ammoniagenesis, redox balance, and calcium signaling among others (reviewed in (45)). Our observations that serum cystatin C, BUN, and tubular injury markers *Ngal* and *Kim1* were normal in uninjured MPC TubKO mice suggest that tubular cell metabolic adaptations to MPC loss are sufficient to maintain normal function and do not overtly cause kidney injury.

Our metabolomic profiling and stable isotope tracing data clearly show that MPC disruption decreases mitochondrial pyruvate oxidation. Both ^13^C-glucose and ^13^C-lactate/pyruvate tracing showed that total kidney ^13^C TCA cycle enrichments were decreased, including M+3 enriched aspartate and citrate, which represent pyruvate anaplerosis that enables fatty acid oxidation. Because the kidney generates the majority of its energy from fatty acid oxidation (46) and normal kidney function was retained after MPC TubKO, our results suggest that essential pyruvate oxidation and anaplerosis were also maintained, likely by a combination of pyruvate-alanine cycling that bypasses the MPC and increased glutaminolysis (18, 27, 28, 31, 41). In addition to TCA cycle adaptations, the increased M+6 fructose-6-phosphate enrichment and pyruvate abundance we observed in MPC TubKO mice is consistent with increased glycolytic flux, which could help sustain ATP production with tubular MPC loss. Previous research has shown that decreased renal gluconeogenesis following kidney injury is associated with a worse prognosis (8, 47), which is consistent with increased glycolysis and decreased mitochondrial pyruvate utilization. Our findings in MPC TubKO mice suggest that that these changes in glucose and pyruvate metabolism are adaptive correlates of but not drivers of AKI severity.

Our results reveal that tubular oxidative state is closely tied to MPC-dependent metabolism. The mitochondrial ETC maintains cellular ATP levels and produces physiologic ROS that informs metabolic regulation. However, perturbations in mitochondrial function can dysregulate ETC function and make it the primary cellular producer of pathological ROS (36, 48). Our observations that uninjured MPC TubKO mice have decreased Complex III activity, increased 3NT-modified proteins, and, in isolated MPC TubKO tubular cells, increased mitochondrial ROS production indicates that tubular MPC loss alters TCA cycle-ETC coupling. This could portend that tubular MPC disruption would be damaging to the kidney, especially during the extreme ROS burden of AKI. Indeed, glutathione depletion has been associated with increased oxidative damage in AKI (49), and delivery of AAV-glutathione reductase has been shown to protect from kidney damage (50). However, the AKI-protective effects from MPC TubKO suggest that the increased antioxidant capacity induced by MPC disruption supersedes the primary metabolic stress imposed by it.

Our results highlight mitochondrial pyruvate oxidation, beyond glycolytic pyruvate production, as contributing to AKI. AKI-dependent deactivating s-nitrosylation of pyruvate kinase M2 (PKM2) was previously found to increase PPP metabolites and protect from ischemia-reperfusion injury (12). However, compared to PKM2, decreasing MPC activity also impairs mitochondrial oxidation of circulating lactate. Lactate is a major kidney fuel source (51), that bypasses glycolysis and is converted to pyruvate before mitochondrial oxidation. Notably, our metabolomic data show the MPC disruption not only impairs pyruvate oxidation but increases kidney lactate levels, which is also observed after AKI (9). Beyond their role as fuels, pyruvate and lactate are chemical antioxidants, and when increased following AKI and in MPC TubKO mice could contribute to the AKI-protective effects we observed in MPC TubKO mice (52-56). In accord, pyruvate administration during AKI has previously been shown to protect against kidney damage (10).

Lastly, we note key limitations of this study and potential future areas of research interest. We do not address the rapid onset mechanism that downregulates *Mpc1* following AKI or the processes increasing glutathione reductase and thioredoxin reductase activities. Understanding these processes could influence novel preemptive approaches to protectively modulate tubular metabolism during AKI. Given the continual discovery and development of MPC inhibitors like MSDC-0602 (57), zaprinast (58), new UK5099-like analogues (59), and non-indole inhibitors (60), modulating MPC activity pharmacologically as a preconditioning strategy or quickly after injury could be a potential approach to minimize AKI-dependent kidney damage. Next, Pax8-dependent *Mpc1* knockout was not complete across all tubular segments. Thus, the stress of completely ablating MPC activity could be greater than we observed. Alternatively, the efficacy of MPC disruption to protect from AKI could be understated.

In conclusion, we show that kidney mitochondrial pyruvate uptake can be modulated to coordinately upregulate oxidant defense systems and protect from AKI. Our data support a model where MPC disruption elicits a goldilocks level of metabolic stress consistent with the concept of hormesis, where a mild, non-injurious stress upregulates stress defense systems, leading to protection from more severe stresses. Given the complexity of the mitochondrial transporter system and its relative lack of direct investigation in the kidney, we expect future research will reveal roles for mitochondrial fuel transport in both exacerbating and protecting from kidney injury and disease.

## METHODS

### Mice breeding

Mice were housed in ventilated cages located in a climate-controlled facility with a 12-hour light/dark cycle. *Ad-libitum* access to water and standard rodent diet was provided, unless otherwise specified. Mice expressing the mitochondrial pyruvate carrier gene flanked by loxP sites (MPC1^f/f^) were generated as previously described (27). The MPC1^f/f^ mice were bred with mice that express Cre recombinase under the control of the tubule specific Pax8^Cre^ promoter (Jackson Laboratory, 028196) (34, 61). Offspring [Pax8^Cre+/-^MPC1^f/f^ (MPC TubKO)] and controls [Pax8^Cre-/-^MPC1^fl/fl^ (WT)] were used for experiments maintained on a C57Bl/6J background. ROSA^mT/mG^ (Jackson Laboratory, 007676) were crossed with mice expressing Cre under the control of the gamma-glutamyltransferase promoter (Ggt1-Cre) to generate mice expressing membrane-localized enhanced green fluorescent (GFP) protein in renal tubular epithelial cells while other cells express membrane-localized tdTomato fluorescent protein as previously described (62).

### Animal models of AKI

Ischemia reperfusion (IR), cisplatin, and rhabdomyolysis induced AKI were induced in 8-12 weeks male mice using methods previously described (63-66). Briefly, to cause ischemia-reperfusion injury, mice were anesthetized by isoflurane and placed on a surgical platform where the body temperature was monitored during the procedure. The skin was disinfected, kidneys were exposed, and bilateral renal pedicles were clamped for 30 minutes, followed by clamp removal, and suturing to close the muscle and skin around the incision. To compensate for the fluid loss, 0.5 ml of warm sterile saline was administered via intraperitoneal injection. The sham groups underwent a similar procedure but without bilateral clamping. Cisplatin induced nephrotoxicity was trigged by a single intraperitoneal cisplatin (30 mg/kg) injection as described previously (65). The sham (vehicle) groups were injected with equal volumes of normal saline. To induce rhabdomyolysis, mice were intramuscularly injected with 50% glycerol (Sigma, G7893, dose 7.5-10 ml/kg) to the two hind-legs or injected with saline as a control.

### Isolation of GFP positive RTECs

mT/mG/Ggt1-Cre mice display renal tubular epithelial cell (RTEC)-specific GFP expression (62, 64). For kidney cell isolation, mice were euthanized, kidneys were excised, and cortical regions were minced and treated with collagenase. Cellular suspensions were passage through 100 μm and 35 μm mesh to generate a single cell population. Anti-GFP antibody and MACS columns (Miltenyi Biotech) were used to isolate GFP-positive RTECs and GFP-negative (tdTomato+) Non-RTECs. The purity (generally greater than 95%) of isolated RTECs and Non-RTECs was verified by flow cytometric analysis.

### RNA isolation and real-time PCR

RNA was isolated from one quarter of a mouse kidney using RNeasy Plus Mini Kit (Qiagen, 74134). For quantitative real-time PCR, an equal amount of RNA was reverse transcribed using Verso cDNA synthesis kit (Fisher Scientific, AB1453B) according to the manufacturer’s protocol. Real-time polymerase chain reaction (PCR) was performed using ABsolute Blue QPCR Mix, SYBR Green (Thermo Scientific, AB4322B). Relative abundance of mRNA was normalized to ribosomal protein U36b4 unless otherwise specified. qRT-PCR primers were designed using Primer-Blast. Primers are listed in **Supplemental Table 1**.

### Western Blot Analysis

Snap-frozen kidney were homogenized using a Tissuelyser in RIPA Lysis and Extraction buffer (Fisher Scientific, PI89901) containing protease inhibitor (Roche, 11836170001) and Phosphatase inhibitor (Millipore, 524625). Crude homogenates were centrifuged at 12000 rpm for 10 minutes and cleared tissues lysates were collected. The protein content of the cleared lysates were quantified using Pierce Rapid Gold BCA Protein Kit (Thermo Scientific, A53226) and separated by Bio-Rad TGX stain-free gel. Separated proteins transferred to PVDF membrane, blocked with PBS supplemented with 5% BSA, and incubated with primary antibodies at 4°C overnight and secondary antibodies for 1 hour. Antibodies are listed in **Supplemental Table 2**.

### Renal Function

Blood samples were centrifuged at 1200 rpm for 2 min at room temperature in BD Microtainer serum separator tubes (BD, 365967) to obtain serum. Serum Cystatin C levels were measured using the Mouse/Rat Cystatin C Quantikine enzyme-linked immunosorbent assay (ELISA) kit (R&D Systems, MSCTC0). Blood Nitrogen Urea serum levels were measured using the QuantiChrom Urea Assay Kit (BioAssay Systems, DIUR-100).

### Blood glucose and lactate measurements

Blood glucose and blood lactate were measured using a One Touch UltraMini glucometer and Nova Biomedical Lactate Plus lactate meter, respectively.

### Immunofluorescence

For histological analysis, kidneys were sliced in half transversely, fixed in 4% paraformaldehyde or formalin for 24 hours, then placed on 70% ethanol until paraffin embedding process was completed and sectioned at 5 μm. After deparaffinized in xylene and rehydrated, antigen retrieval was performed using 10 mM sodium citrate, pH 6.0, with 0.05% Tween for 25 min in pressure cooker. Sections were washed in 1X PBS 0.05% Tween, blocked using Super Block (ScyTek Laboratories, AAA125) 10 minutes and incubated with primary antibodies diluted in Normal Antibody Diluent (ScyTek Laboratories, ABB125) at 4°C overnight, and secondary antibodies for 1 hour according (**Supplemental Table 2**). Sections were then mounted using VECTASHIELD Vibrance Antifade Mounting Medium with DAPI (Vector Laboratories, H-1800), or Prolong Gold (Life Technologies, P36931). Images were obtained using an Olympus BX51 microscope and CX9000 camera (Shinjuku, Japan), or by confocal imaging using Olympus FV1000 confocal laser scanning microscope (Olympus America).

### Immunofluorescent and IHC measures of oxidation

Paraffin-embedded kidneys were sectioned at 4 μm for evaluation of 3-nitrotyrosine (3-NT) and protein-glutathionylation (supplemental methods).

### Tubular injury score

For histological analysis, paraffin-embedded kidneys were sectioned at 5 μm and stained with Period Acid-Shift (PAS) for kidney injury semiquantitative evaluation (67, 68) by a blinded pathologist (supplemental methods).

### Tunel assay

Apoptotic cell death was detected using the The ApopTag Plus Fluorescein In Situ Apoptosis Detection Kit (Sigma, S7111, supplemental methods).

### Mitochondrial and redox response evaluation

Mitochondrial electron transport chain activity assays, TCA cycle activity assays, total glutathione quantification, oxidized glutathione quantification, and antioxidant response activity assays performed in frozen kidney tissue (supplemental methods).

### MitoSOX Oxidation in Primary Culture Renal Tubule Epithelial Cells

Primary Tubular Cells were obtained from 8 - 10-week-old MPC TubKO and littermate WT control mice (supplemental methods). Primary Tubular Cells were grown in 60 mm cell culture dishes. After two days, cells were washed with ice-cold DPBS and trypsinized for 3 minutes at 37°C, washed with PBS without Ca/Mg containing 5 mM pyruvate, and centrifuged at 1200 rpm for 5 minutes. Cells were incubated with 2 μM MitoSOX Red Mitochondrial Superoxide Indicator (Invitrogen, M36008) for 15 minutes at 37°C in dark. 10 μM of Antimycin A (Sigma Aldrich, A8674) was used as a positive control. To stop the reaction, cells were placed on ice and then transferred into Falcon tubes (Falcon, 352235). Samples were analyzed using a BD LSRII Flow Cytometer. The mean fluorescence intensity (MFI) of 10,000 events was analyzed in each sample and corrected for autofluorescence from unlabeled cells using FlowJo software (Tree Star, Ashland, OR). The MFI data were normalized to WT control.

### Mitochondrial isolation and oxygen consumption rate (OCR) measurements

Mitochondria were isolated by differential centrifugation (supplemental methods). To measure mitochondrial oxygen consumption, mitochondrial pellets were resuspended in a buffer containing 70 mM Sucrose, 220 mM d-mannitol, 10 mM KH2PO4, 5 mM MgCl2, 5 mM HEPES pH 7.2, 1 mM EGTA, and 0.2% fatty acid free BSA. Mitochondrial protein content of the suspensions was determined by Bradford Assay and 5 μg of kidney mitochondria were attached per well of the V3-PET seahorse plates by centrifugation at 2000 x g for 20 minutes. Substrate-containing buffer was added such that the final concentrations were 15 mM Glutamate and 1 mM Malate. A Seahorse assay was performed where cycles of a 1-minute mix step, a 1-minute wait step, and a 3-minute measurement step were conducted. Three cycles of basal measurements were acquired before three cycles of stimulated respiration was measured following the addition of 4 mM ADP and 1 μM FCCP. Finally, three cycles of ETC-inhibited respiration were measured following the addition of 5 μM rotenone. Oxygen consumption was normalized to protein loading, and three measurement cycles per state (basal, stimulated, and ETC-inhibited) were averaged and normalized to the basal oxygen consumption rate.

### Metabolomics and data analysis

Whole kidneys were rapidly dissected from live mice under isoflurane anesthesia and then within 1-2 seconds freeze clamped with liquid nitrogen temperature tongs. Kidneys were extracted for metabolomics analysis (supplemental methods) by gas chromatography (GC)- and liquid chromatography (LC)-mass spectrometry (MS) as previously described (37). Acquired MS data were processed by Thermo Scientific TraceFinder 4.1 software. Metabolites were identified by matching with the University of Iowa Metabolomics Core Facility standard-confirmed, in-house libraries documenting a retention time, a target ion, and at least 1 confirming ion per metabolite (GC) or retention time, accurate mass, and MS/MS data when available (LC). The NOREVA tool was used to correct for instrument drift by regressing peak intensities from experimental samples against those of pooled QC samples analyzed throughout the run (69). NOREVA corrected data were normalized to the D4-succinate signal/sample to control for extraction, derivatization (GC), and/or loading effects.

### ^13^C-glucose and ^13^C-lactate/^13^C-pyruvate in vivo tracing

7 - 8- or 10 - 12-week-old Mpc1^f/f^ and Mpc1^f/f^Pax8^+^ mice were fasted for 4 hours prior to intraperitoneal. injection with ^13^C-glucose (10% ^13^ C-glucose, 2.0g/kg lean body mass, Cambridge Isotope) or ^13^C-lactate/^13^C-pyruvate (10:1, 3.0g/kg lean body mass, Cambridge Isotope), respectively. 30 minutes after injection mice kidneys were collected from isoflurane anesthetized mice by freeze-clamping as described above. Freeze-clamped tissues were processed as described above. Kidneys harvested from mice treated with natural abundance lactate/pyruvate was used to correct for ^13^C natural abundance (70). Metabolites were identified as described above.

### Statistics

Data are presented as the mean ± SEM. Statistical analysis was performed using GraphPad Prism 8 and 9 (GraphPad Software). An unpaired t test with Welch’s correction was used to compare differences between 2 groups. Multiple-group comparisons were performed using one- or two-way ANOVA with Tukey’s multiple comparison test. Differences were considered statistically significant when a P value was less than 0.05.

### Study approval

All animal care and experimental procedures were in adherence of the National Institute of Health Guide for the Care and Use of Laboratory Animals and Institutional Committee policies. Studies were approved by the Nationwide Children’s Hospital Institutional Animal Care and Use Committee protocol #AR20-00055 and by the University of Iowa Institutional Animal Care and Use Committee protocol #8041235-004.

## DATA AVAILABILITY

All relevant data supporting the key findings of this study are available within the article and its Supplementary Information files or from the corresponding authors upon reasonable request.

## ACKNOWLEDGEMENTS

This work was supported by grants CHD K12 HD027748 (DZO), NIH R01 DK104998 and the University of Iowa Healthcare Distinguished Scholars Award (EBT), ADA 1-18-PDF-060 and AHA CDA851976 (AJR), NIH P01 CA217797 and P30 CA086862 (DRS, BGA, KAM, MLM), NIDDK K01 DK126991 (ARJ), T32 DK007690 (EJS), and NIAMS R00 AR070914 (MCC).

## AUTHOR CONTRIBUTIONS

EBT and DZO conceived the study. AJR, GVM, DRS, EBT, and DZO designed the study. AJR, GVM, GMA, HW, JYK, AS, KM, PR, EJS, AJ, and MLM performed experiments and collected data. AJR, GVM, DRS, EBT, and DZO analyzed data. AJR, GVM, BA, NP, ARJ, MCC, DRS, EBT, and DZO interpreted data. AJR, GVM, DRS, EBT, and DZO wrote the draft manuscript. All authors read, revised, and approve the final manuscript. EBT and DZO supervised the study.AJR and GVM are co-first authors. The order of the cofirst authors was determined based on their efforts and contributions to the manuscript.

## REFERENCES

1. Chawla LS, Bellomo R, Bihorac A, Goldstein SL, Siew ED, Bagshaw SM, et al. Acute kidney disease and renal recovery: consensus report of the Acute Disease Quality Initiative (ADQI) 16 Workgroup. Nat Rev Nephrol. 2017;13(4):241–57.

2. Chertow GM, Burdick E, Honour M, Bonventre JV, and Bates DW. Acute kidney injury, mortality, length of stay, and costs in hospitalized patients. J Am Soc Nephrol. 2005;16(11):3365–70.

3. Coca SG, Singanamala S, and Parikh CR. Chronic kidney disease after acute kidney injury: a systematic review and meta-analysis. Kidney Int. 2012;81(5):442–8.

4. Goldstein SL, Jaber BL, Faubel S, Chawla LS, and Acute Kidney Injury Advisory Group of American Society of N. AKI transition of care: a potential opportunity to detect and prevent CKD. Clin J Am Soc Nephrol. 2013;8(3):476–83.

5. Bhargava P, and Schnellmann RG. Mitochondrial energetics in the kidney. Nat Rev Nephrol. 2017;13(10):629–46.

6. Kaddourah A, Basu RK, Bagshaw SM, Goldstein SL, and Investigators A. Epidemiology of Acute Kidney Injury in Critically Ill Children and Young Adults. N Engl J Med. 2017;376(1):11–20.

7. Grgic I, Campanholle G, Bijol V, Wang C, Sabbisetti VS, Ichimura T, et al. Targeted proximal tubule injury triggers interstitial fibrosis and glomerulosclerosis. Kidney Int. 2012;82(2):172–83.

8. Legouis D, Ricksten SE, Faivre A, Verissimo T, Gariani K, Verney C, et al. Altered proximal tubular cell glucose metabolism during acute kidney injury is associated with mortality. Nat Metab. 2020;2(8):732–43.

9. Lan R, Geng H, Singha PK, Saikumar P, Bottinger EP, Weinberg JM, et al. Mitochondrial Pathology and Glycolytic Shift during Proximal Tubule Atrophy after Ischemic AKI. J Am Soc Nephrol. 2016;27(11):3356–67.

10. Zager RA, Johnson AC, and Becker K. Renal cortical pyruvate depletion during AKI. J Am Soc Nephrol. 2014;25(5):998–1012.

11. Shen Y, Jiang L, Wen P, Ye Y, Zhang Y, Ding H, et al. Tubule-derived lactate is required for fibroblast activation in acute kidney injury. Am J Physiol Renal Physiol. 2020;318(3):F689–F701.

12. Zhou HL, Zhang R, Anand P, Stomberski CT, Qian Z, Hausladen A, et al. Metabolic reprogramming by the S-nitroso-CoA reductase system protects against kidney injury. Nature. 2019;565(7737):96–100.

13. Scholz H, Boivin FJ, Schmidt-Ott KM, Bachmann S, Eckardt KU, Scholl UI, et al. Kidney physiology and susceptibility to acute kidney injury: implications for renoprotection. Nat Rev Nephrol. 2021;17(5):335–49.

14. Jang C, Chen L, and Rabinowitz JD. Metabolomics and Isotope Tracing. Cell. 2018;173(4):822–37.

15. Bricker DK, Taylor EB, Schell JC, Orsak T, Boutron A, Chen YC, et al. A mitochondrial pyruvate carrier required for pyruvate uptake in yeast, Drosophila, and humans. Science. 2012;337(6090):96–100.

16. Herzig S, Raemy E, Montessuit S, Veuthey JL, Zamboni N, Westermann B, et al. Identification and functional expression of the mitochondrial pyruvate carrier. Science. 2012;337(6090):93–6.

17. Vacanti NM, Divakaruni AS, Green CR, Parker SJ, Henry RR, Ciaraldi TP, et al. Regulation of substrate utilization by the mitochondrial pyruvate carrier. Mol Cell. 2014;56(3):425–35.

18. Yang C, Ko B, Hensley CT, Jiang L, Wasti AT, Kim J, et al. Glutamine oxidation maintains the TCA cycle and cell survival during impaired mitochondrial pyruvate transport. Mol Cell. 2014;56(3):414–24.

19. Schell JC, Wisidagama DR, Bensard C, Zhao H, Wei P, Tanner J, et al. Control of intestinal stem cell function and proliferation by mitochondrial pyruvate metabolism. Nat Cell Biol. 2017;19(9):1027–36.

20. Grenell A, Wang Y, Yam M, Swarup A, Dilan TL, Hauer A, et al. Loss of MPC1 reprograms retinal metabolism to impair visual function. Proc Natl Acad Sci U S A. 2019;116(9):3530–5.

21. Fernandez-Caggiano M, Kamynina A, Francois AA, Prysyazhna O, Eykyn TR, Krasemann S, et al. Mitochondrial pyruvate carrier abundance mediates pathological cardiac hypertrophy. Nat Metab. 2020;2(11):1223–31.

22. McCommis KS, Kovacs A, Weinheimer CJ, Shew TM, Koves TR, Ilkayeva OR, et al. Nutritional modulation of heart failure in mitochondrial pyruvate carrier-deficient mice. Nat Metab. 2020;2(11):1232–47.

23. Zhang Y, Taufalele PV, Cochran JD, Robillard-Frayne I, Marx JM, Soto J, et al. Mitochondrial pyruvate carriers are required for myocardial stress adaptation. Nat Metab. 2020;2(11):1248–64.

24. Cluntun AA, Badolia R, Lettlova S, Parnell KM, Shankar TS, Diakos NA, et al. The pyruvate-lactate axis modulates cardiac hypertrophy and heart failure. Cell Metab. 2021;33(3):629–48 e10.

25. Schell JC, Olson KA, Jiang L, Hawkins AJ, Van Vranken JG, Xie J, et al. A role for the mitochondrial pyruvate carrier as a repressor of the Warburg effect and colon cancer cell growth. Mol Cell. 2014;56(3):400–13.

26. Compan V, Pierredon S, Vanderperre B, Krznar P, Marchiq I, Zamboni N, et al. Monitoring Mitochondrial Pyruvate Carrier Activity in Real Time Using a BRET-Based Biosensor: Investigation of the Warburg Effect. Mol Cell. 2015;59(3):491–501.

27. Gray LR, Sultana MR, Rauckhorst AJ, Oonthonpan L, Tompkins SC, Sharma A, et al. Hepatic Mitochondrial Pyruvate Carrier 1 Is Required for Efficient Regulation of Gluconeogenesis and Whole-Body Glucose Homeostasis. Cell Metab. 2015;22(4):669–81.

28. McCommis KS, Chen Z, Fu X, McDonald WG, Colca JR, Kletzien RF, et al. Loss of Mitochondrial Pyruvate Carrier 2 in the Liver Leads to Defects in Gluconeogenesis and Compensation via Pyruvate-Alanine Cycling. Cell Metab. 2015;22(4):682–94.

29. Rauckhorst AJ, Gray LR, Sheldon RD, Fu X, Pewa AD, Feddersen CR, et al. The mitochondrial pyruvate carrier mediates high fat diet-induced increases in hepatic TCA cycle capacity. Mol Metab. 2017;6(11):1468–79.

30. Sharma A, Oonthonpan L, Sheldon RD, Rauckhorst AJ, Zhu Z, Tompkins SC, et al. Impaired skeletal muscle mitochondrial pyruvate uptake rewires glucose metabolism to drive whole-body leanness. Elife. 2019;8.

31. Tompkins SC, Sheldon RD, Rauckhorst AJ, Noterman MF, Solst SR, Buchanan JL, et al. Disrupting Mitochondrial Pyruvate Uptake Directs Glutamine into the TCA Cycle away from Glutathione Synthesis and Impairs Hepatocellular Tumorigenesis. Cell Rep. 2019;28(10):2608–19 e6.

32. Kirita Y, Wu H, Uchimura K, Wilson PC, and Humphreys BD. Cell profiling of mouse acute kidney injury reveals conserved cellular responses to injury. Proc Natl Acad Sci U S A. 2020;117(27):15874–83.

33. Kim JY, Bai Y, Jayne LA, Abdulkader F, Gandhi M, Perreau T, et al. SOX9 promotes stress-responsive transcription of VGF nerve growth factor inducible gene in renal tubular epithelial cells. J Biol Chem. 2020;295(48):16328–41.

34. Bouchard M, Souabni A, Mandler M, Neubuser A, and Busslinger M. Nephric lineage specification by Pax2 and Pax8. Genes Dev. 2002;16(22):2958–70.

35. Nizar JM, Shepard BD, Vo VT, and Bhalla V. Renal tubule insulin receptor modestly promotes elevated blood pressure and markedly stimulates glucose reabsorption. JCI Insight. 2018;3(16).

36. Chouchani ET, Pell VR, Gaude E, Aksentijevic D, Sundier SY, Robb EL, et al. Ischaemic accumulation of succinate controls reperfusion injury through mitochondrial ROS. Nature. 2014;515(7527):431–5.

37. Rauckhorst AJ, Borcherding N, Pape DJ, Kraus AS, Scerbo DA, and Taylor EB. Mouse tissue harvest-induced hypoxia rapidly alters the in vivo metabolome, between-genotype metabolite level differences, and (13)C-tracing enrichments. Mol Metab. 2022;66:101596.

38. Goodman RP, Markhard AL, Shah H, Sharma R, Skinner OS, Clish CB, et al. Hepatic NADH reductive stress underlies common variation in metabolic traits. Nature. 2020;583(7814):122–6.

39. Thompson Legault J, Strittmatter L, Tardif J, Sharma R, Tremblay-Vaillancourt V, Aubut C, et al. A Metabolic Signature of Mitochondrial Dysfunction Revealed through a Monogenic Form of Leigh Syndrome. Cell Rep. 2015;13(5):981–9.

40. Lu SC. Glutathione synthesis. Biochim Biophys Acta. 2013;1830(5):3143–53.

41. Bowman CE, Zhao L, Hartung T, and Wolfgang MJ. Requirement for the Mitochondrial Pyruvate Carrier in Mammalian Development Revealed by a Hypomorphic Allelic Series. Mol Cell Biol. 2016;36(15):2089–104.

42. Petejova N, and Martinek A. Acute kidney injury due to rhabdomyolysis and renal replacement therapy: a critical review. Crit Care. 2014;18(3):224.

43. Giustarini D, Dalle-Donne I, Milzani A, Braconi D, Santucci A, and Rossi R. Membrane Skeletal Protein S-Glutathionylation in Human Red Blood Cells as Index of Oxidative Stress. Chem Res Toxicol. 2019;32(6):1096–102.

44. Schafer FQ, and Buettner GR. Redox environment of the cell as viewed through the redox state of the glutathione disulfide/glutathione couple. Free Radic Biol Med. 2001;30(11):1191–212.

45. Gewin LS. Sugar or Fat? Renal Tubular Metabolism Reviewed in Health and Disease. Nutrients. 2021;13(5).

46. Nieth H, and Schollmeyer P. Substrate-utilization of the human kidney. Nature. 1966;209(5029):1244–5.

47. Verissimo T, Faivre A, Rinaldi A, Lindenmeyer M, Delitsikou V, Veyrat-Durebex C, et al. Decreased Renal Gluconeogenesis Is a Hallmark of Chronic Kidney Disease. J Am Soc Nephrol. 2022;33(4):810–27.

48. Liu Y, Fiskum G, and Schubert D. Generation of reactive oxygen species by the mitochondrial electron transport chain. J Neurochem. 2002;80(5):780–7.

49. Shang Y, Siow YL, Isaak CK, and O K. Downregulation of Glutathione Biosynthesis Contributes to Oxidative Stress and Liver Dysfunction in Acute Kidney Injury. Oxid Med Cell Longev. 2016;2016:9707292.

50. Gao D, Wang S, Lin Y, and Sun Z. In vivo AAV delivery of glutathione reductase gene attenuates anti-aging gene klotho deficiency-induced kidney damage. Redox Biol. 2020;37:101692.

51. Hui S, Ghergurovich JM, Morscher RJ, Jang C, Teng X, Lu W, et al. Glucose feeds the TCA cycle via circulating lactate. Nature. 2017;551(7678):115–8.

52. Groussard C, Morel I, Chevanne M, Monnier M, Cillard J, and Delamarche A. Free radical scavenging and antioxidant effects of lactate ion: an in vitro study. J Appl Physiol (1985). 2000;89(1):169–75.

53. Guarino VA, Oldham WM, Loscalzo J, and Zhang YY. Reaction rate of pyruvate and hydrogen peroxide: assessing antioxidant capacity of pyruvate under biological conditions. Sci Rep. 2019;9(1):19568.

54. Tauffenberger A, Fiumelli H, Almustafa S, and Magistretti PJ. Lactate and pyruvate promote oxidative stress resistance through hormetic ROS signaling. Cell Death Dis. 2019;10(9):653.

55. Ramos-Ibeas P, Barandalla M, Colleoni S, and Lazzari G. Pyruvate antioxidant roles in human fibroblasts and embryonic stem cells. Mol Cell Biochem. 2017;429(1-2):137–50.

56. Wang X, Perez E, Liu R, Yan LJ, Mallet RT, and Yang SH. Pyruvate protects mitochondria from oxidative stress in human neuroblastoma SK-N-SH cells. Brain Res. 2007;1132(1):1–9.

57. Vigueira PA, McCommis KS, Hodges WT, Schweitzer GG, Cole SL, Oonthonpan L, et al. The beneficial metabolic effects of insulin sensitizers are not attenuated by mitochondrial pyruvate carrier 2 hypomorphism. Exp Physiol. 2017;102(8):985–99.

58. Du J, Cleghorn WM, Contreras L, Lindsay K, Rountree AM, Chertov AO, et al. Inhibition of mitochondrial pyruvate transport by zaprinast causes massive accumulation of aspartate at the expense of glutamate in the retina. J Biol Chem. 2013;288(50):36129–40.

59. Hegazy L, Gill LE, Pyles KD, Kaiho C, Kchouk S, Finck BN, et al. Identification of Novel Mitochondrial Pyruvate Carrier Inhibitors by Homology Modeling and Pharmacophore-Based Virtual Screening. Biomedicines. 2022;10(2).

60. Liu X, Flores AA, Situ L, Gu W, Ding H, Christofk HR, et al. Development of Novel Mitochondrial Pyruvate Carrier Inhibitors to Treat Hair Loss. J Med Chem. 2021;64(4):2046–63.

61. Grouls S, Iglesias DM, Wentzensen N, Moeller MJ, Bouchard M, Kemler R, et al. Lineage specification of parietal epithelial cells requires beta-catenin/Wnt signaling. J Am Soc Nephrol. 2012;23(1):63–72.

62. Bai Y, Kim JY, Bisunke B, Jayne LA, Silvaroli JA, Balzer MS, et al. Kidney toxicity of the BRAF-kinase inhibitor vemurafenib is driven by off-target ferrochelatase inhibition. Kidney Int. 2021;100(6):1214–26.

63. Kim JY, Bai Y, Jayne LA, Cianciolo RE, Bajwa A, and Pabla NS. Involvement of the CDKL5-SOX9 signaling axis in rhabdomyolysis-associated acute kidney injury. Am J Physiol Renal Physiol. 2020;319(5):F920–F9.

64. Kim JY, Bai Y, Jayne LA, Hector RD, Persaud AK, Ong SS, et al. A kinome-wide screen identifies a CDKL5-SOX9 regulatory axis in epithelial cell death and kidney injury. Nat Commun. 2020;11(1):1924.

65. Kim JY, Jayne LA, Bai Y, Feng M, Clark MA, Chung S, et al. Ribociclib mitigates cisplatin-associated kidney injury through retinoblastoma-1 dependent mechanisms. Biochem Pharmacol. 2020;177:113939.

66. Pabla N, Gibson AA, Buege M, Ong SS, Li L, Hu S, et al. Mitigation of acute kidney injury by cell-cycle inhibitors that suppress both CDK4/6 and OCT2 functions. Proc Natl Acad Sci U S A. 2015;112(16):5231–6.

67. Hur E, Garip A, Camyar A, Ilgun S, Ozisik M, Tuna S, et al. The effects of vitamin d on gentamicin-induced acute kidney injury in experimental rat model. Int J Endocrinol. 2013;2013:313528.

68. Zhou XJ, Laszik Z, Wang XQ, Silva FG, and Vaziri ND. Association of renal injury with increased oxygen free radical activity and altered nitric oxide metabolism in chronic experimental hemosiderosis. Lab Invest. 2000;80(12):1905–14.

69. Li B, Tang J, Yang Q, Li S, Cui X, Li Y, et al. NOREVA: normalization and evaluation of MS-based metabolomics data. Nucleic Acids Res. 2017;45(W1):W162–W70.

70. Fernandez CA, Des Rosiers C, Previs SF, David F, and Brunengraber H. Correction of 13C mass isotopomer distributions for natural stable isotope abundance. J Mass Spectrom. 1996;31(3):255–62.

