## Supplemental Document for "Tubular Mitochondrial Pyruvate Carrier Disruption Elicits Redox Adaptations that Protect from Acute Kidney Injury"

### Tubular Mitochondrial Pyruvate Carrier Disruption Results in Redox Adaptation and Protects from Acute Kidney Injury

**SUPPLEMENTAL TABLE 1:** qPCR primers

**SUPPLEMENTAL TABLE 2:** Western blotting and immunostaining antibodies

**SUPPLEMENTAL TABLE 3:** Metabolomic profiling data in WT and MPC TubKO kidneys

**SUPPLEMENTAL TABLE 4:** $^{13}$C-glucose isotoplogue enrichment and abundance data for WT and MPC TubKO kidneys

**SUPPLEMENTAL TABLE 5:** $^{13}$C-lactate and -pyruvate isotoplogue enrichment and abundance data for WT and MPC TubKO kidneys

**SUPPLEMENTAL TABLE 6:** Tubular injury score criteria

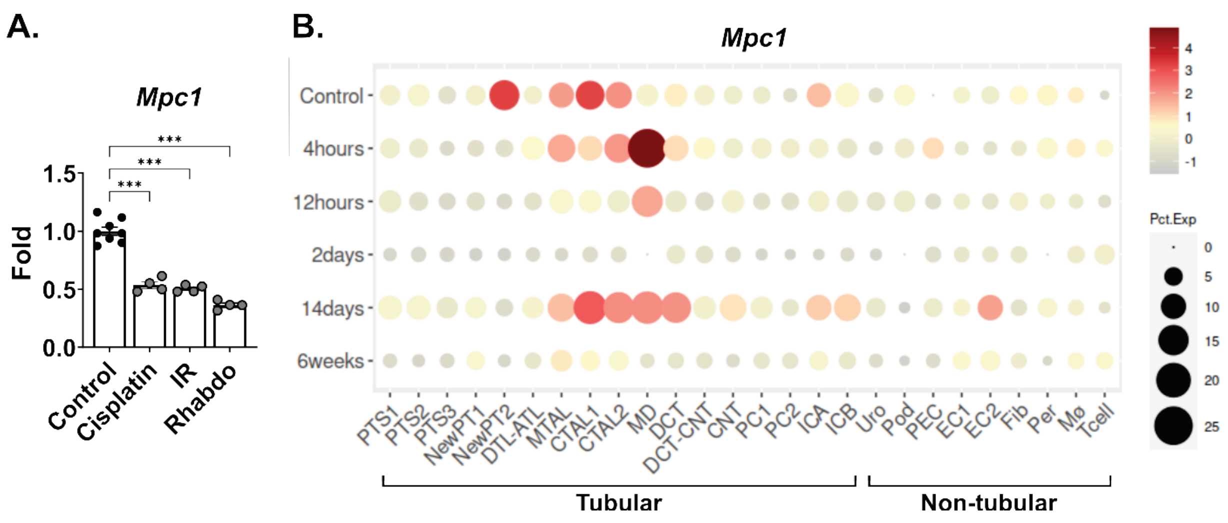


**SUPPLEMENTAL FIGURE 1. *Mpc1* RNA is expressed in all tubular segments and is disrupted in AKI.** (**A**) Bar graph showing *Mpc1* mRNA level in different kidney injury models obtained from a previously published RNAseq dataset (1). (n = 8/group for control, n = 4/group for injury, *** p < 0.001 by one-way ANOVA with Dunnett's multiple comparison test).
(**B**) Single cell heatmap of *Mpc1* levels in kidneys obtained at different time points following ischemia reperfusion-induced AKI. Data visualization was performed using the interactive transcriptomics online analyzer at www.Humphreyslab.com/SingleCell/ (2).
Data are presented as means ± SEM.

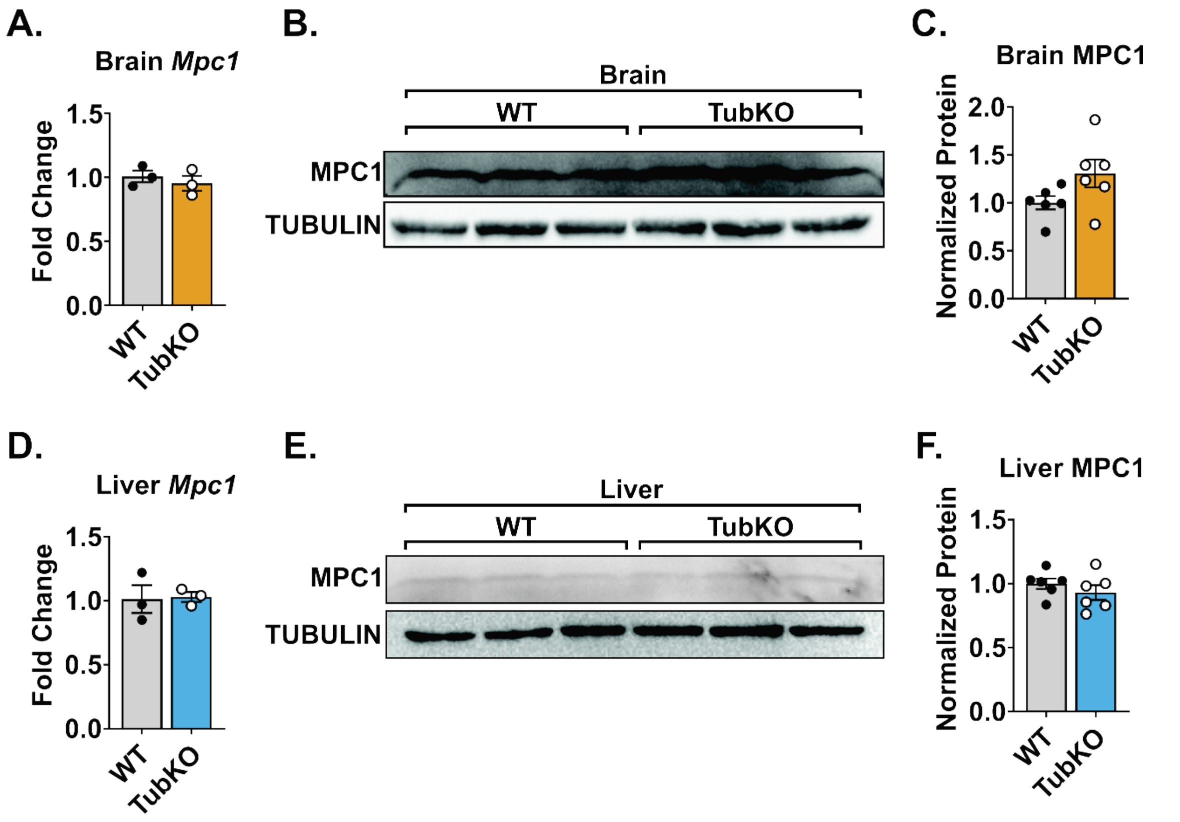


**SUPPLEMENTAL FIGURE 2. Confirmation of selective *Mpc1* deletion in MPC TubKO mice.**
(**A**) Bar graph showing brain *Mpc1* mRNA levels in WT and MPC TubKO mice. (n = 3/group, 8 - 12-week-old mice).
(**B-C**) Representative Western blot (**B**) and quantification (**C**) of brain MPC1 protein abundance in WT and MPC TubKO mice. TUBULIN was blotted as loading control and used as the protein quantification normalizer. (n = 3/group for Western blot and n = 6/group for qPCR, 8 - 12-week-old mice).
(**D**) Bar graph showing liver *Mpc1* mRNA levels in WT and MPC TubKO mice. (n = 3/group, 8 - 12-week-old mice).
(**E-F**) Representative Western blot (**E**) and quantification (**F**) of liver MPC1 protein abundance in WT and MPC TubKO mice. TUBULIN was blotted as loading control and used as the protein quantification normalizer. (n = 3/group for Western blot and n = 6/group for qPCR, 8 - 12-week-old mice).
Data are presented as means ± SEM.

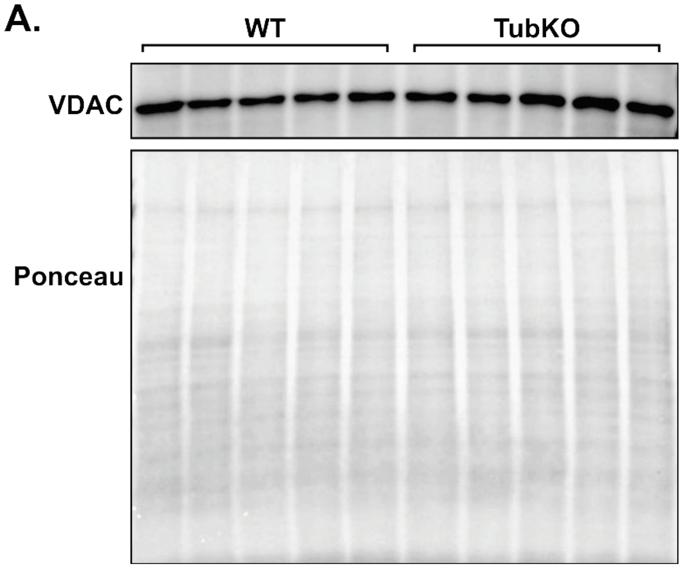


**SUPPLEMENTAL FIGURE 3. Kidney VDAC protein abundance is unchanged in MPC TubKO mice.**
(**A**) Representative Western blot of kidney VDAC protein abundance and a ponceau-stained gel image depicting total protein content of kidney lysates from MPC TubKO mice. (n = 5/group, 6 - 8-week-old mice).

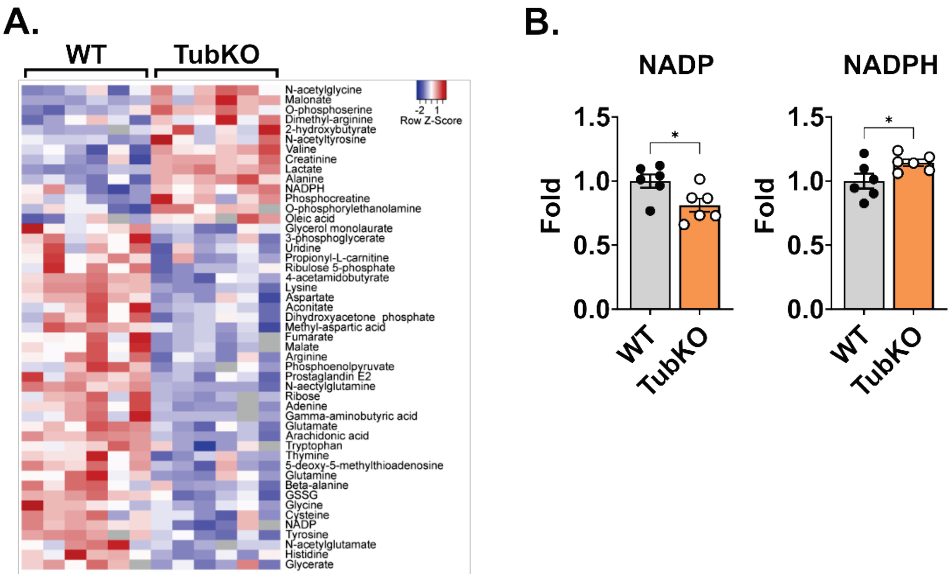


**SUPPLEMENTAL FIGURE 4. MPC TubKO mice have mitochondrial redox adaptation in vivo.**
(**A**) Heat map showing significantly different kidney metabolites in WT and MPC TubKO mice. (n = 6/group, 8 - 12-week-old mice, $p < 0.05$, by unpaired t test correction).
(**B**) Bar graphs show kidney NADP and NADPH levels in WT and MPC TubKO mice. (n = 6/group, 8 - 12-week-old mice, * $p < 0.05$, by unpaired t test correction).
Data are presented as means $\pm$ SEM.

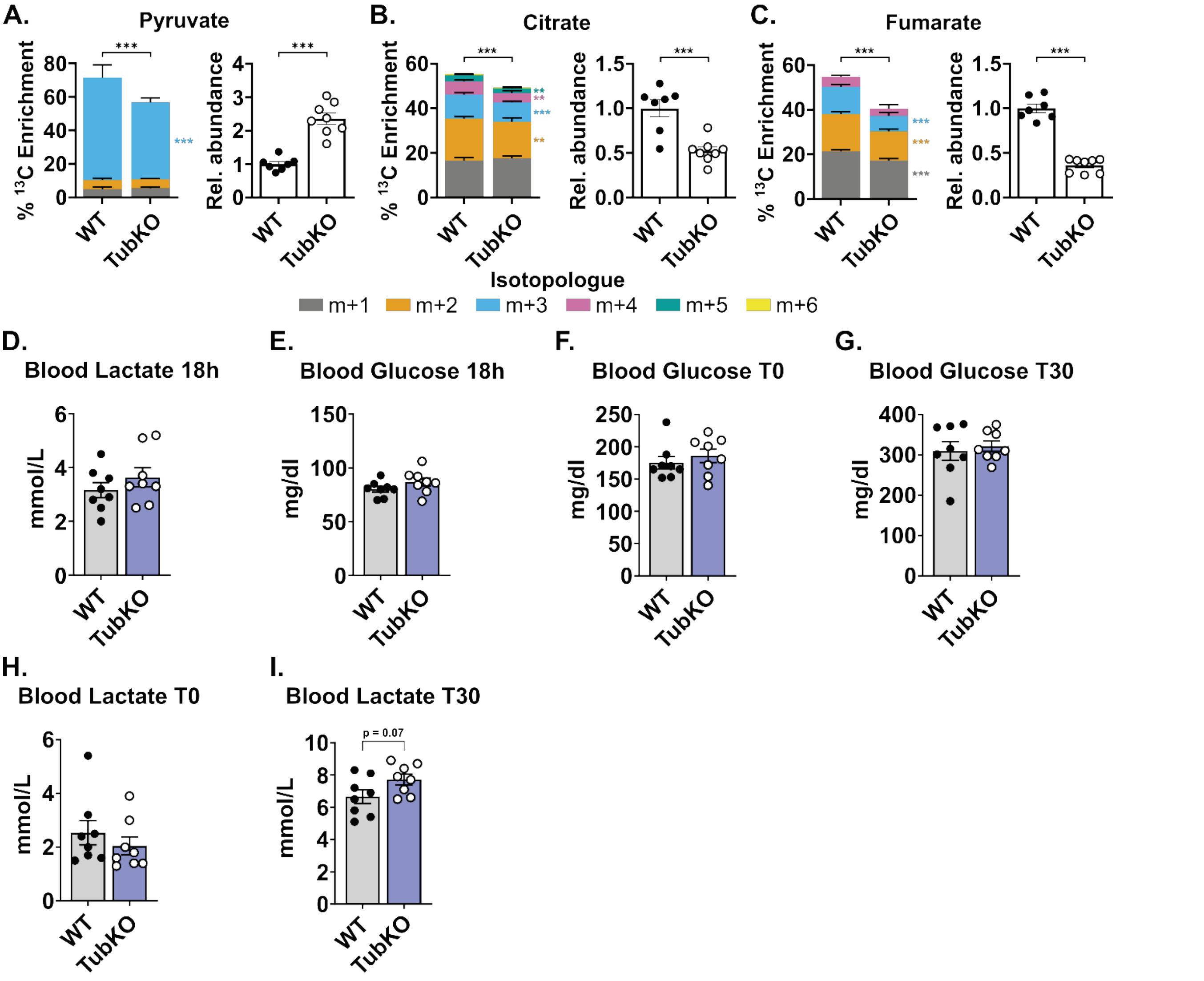


**SUPPLEMENTAL FIGURE 5. MPC TubKO mice have decreased mitochondrial pyruvate oxidation that does not affect systemic glucose homeostasis.**

**(A-C)** Stacked bar graphs showing kidney $^{13}C$-isotopologue enrichments and bar graphs relative abundances of metabolites 30 minutes after $^{13}C$-lactate/$^{13}C$-pyruvate bolus injection in WT and MPC TubKO mice. Pyruvate (**A**), citrate (**B**), and fumarate (**C**). (n = 7 - 8/group, 10 - 12-week-old mice, ** $p < 0.01$, *** $p < 0.001$ by unpaired t test).

(**D-E**) Bar graphs showing systemic blood lactate (**A**) and glucose (**B**) levels after an 18 hour fast in WT and MPC TubKO mice. (n = 8, 9 - 11-week-old mice).

(**F-I**) Bar graphs showing systemic blood glucose (**E, F**) and lactate (**H, I**) levels before (**E, H**) and 30 minutes after $^{13}C$-lactate/$^{13}C$-pyruvate bolus injection (**F, I**) in WT and MPC TubKO mice. (n = 8/group, 10 - 12-week-old mice, p - value by unpaired t test with Welch's correction).

Data are presented as means $\pm$ SEM.

**SUPPLEMENTAL METHODS**

**Immunofluorescent measures of tubular protein-glutathionylation.** Paraffin-embedded kidneys were sectioned at 4 µm for evaluation. Immunofluorescence staining were performed using on a Discovery Ultra autostaining system (Roche). Primary antibody staining for glutathione (Abcam, ab19534) was visualized using secondary Omnimap horseradish peroxidase (Roche, 05269652001) followed by Discovery Cy5. Nuclei were stained with Quantum Dot DAPI (Roche). Slides were coverslipped and scanned using a 20X objective at 405 nm and 633 nm with a VS200 scanning microscope (Olympus). Images were analyzed by ImageJ software by blinded selection and quantification of tubular GSH expression in 3 different images taken at 10x magnification. GSH tubular expression was normalized to vehicle control.

**3-nitrotyrosine (3-NT) IHC detection**. 3 µm tissue sections were deparaffinized and rehydrated, followed by antigen retrieval, and blocking using 0.1% fish skin gelatin (Sigma, G7765) and 5% normal horse serum. The primary antibody, 3-NT, was incubated overnight at 4$^{o}$C, followed by secondary antibody, and DAB staining (**Supplemental Table 2**). Three representative images of each kidney section were captured by the Olympus BX-61 Light Microscope. Images were analyzed by ImageJ software for the percent positive DAB stained area (3). The average of the three images per kidney section were reported.

**Tubular injury score.** Tubular necrosis was defined as loss of epithelial cells of the nucleus, dark acidophilic cytoplasm, loss of tubular epithelial cells into tubular lumen, and acellular sections of tubules and it was scored from 0 to 4 according to percentage involvement (**Supplemental Table 6**). Representative figures were taken at 20X magnification using an Olympus BX61 Light Microscope (Olympus).

**Tunel assay.** Apoptotic cell death was detected using the The ApopTag Plus Fluorescein In Situ Apoptosis Detection Kit (Sigma, S7111). Briefly, formalin-fixed, paraffin-embedded kidneys were sectioned at 5 µm. Slides were deparaffinized in xylene and rehydrated. To improve the exposure of DNA by digesting DNA-binding proteins, heat induced epitope retrieval was performed using sodium citrate buffer composed of 10 mM sodium citrate, pH 6.0 with 0.05% Tween 20. Equilibration buffer was applied for 2 minutes, tdT enzyme solution was applied for 1 hour, stop/wash solution was applied for 10 minutes, and anti-digoxigenin conjugate was applied for 30 minutes. Slides were cover-slipped using Vectashield Vibrance Antifade Mounting Medium with DAPI (Vector Laboratories, H-1800), and 3-5 20X magnification images were taken per slide using an Olympus BX51 Light Microscope (Olympus). To score apoptosis, a blinded individual counted TUNEL+ tubular cells in a 250x250 grid and a score was assigned (0: None, 1: 1-5 TUNEL+ cells, 2: 6-10 TUNEL+ cells, 3: >10 TUNEL+ cells). Then, a final image score was calculated with the sum of each image grid's score. The average score of 3-5 images was calculated per mouse.

**Glutathione assay.** Kidney tissue was homogenized on 5% 5-Sulfosalicylic acid dihydrate (Thermo Scientific, AA4314414). Homogenates were centrifuged to remove precipitated proteins and supernatants were collected. The protein pellets were stored and later resuspended in 0.1 N NaOH for protein quantification using the BCA Assay Kit (Thermo Scientific, 23225) for normalization. Total glutathione (GSH) content was determined in the supernatants by the method of Griffith (4). Glutathione disulfide (GSSG) was measured in the supernatant by adding 20 µl of a 1:1 mixture of 2-vinylpyridine and ethanol per 100 µl of sample and incubating for 2 h prior to assaying as described previously (5).

**Activity assays.** For all enzymatic activity assays, kidney tissue was homogenized in DETAPAC buffer comprising 0.05 M phosphate buffer, pH 7.8, 1.34 mmol/L DETAPAC, and protein levels were determined using the Lowry assay. The assays were performed on a Beckman DU 800 spectrophotometer.

*MnSOD activity assay*. SOD activity was determined using the indirect competitive inhibition assay as described previously (6). Briefly, the flux of superoxide generated by the reaction of xanthine & xanthine oxidase was measured using nitroblue tetrazolium (NBT). Increasing concentrations of SOD rapidly converts the superoxide produced to hydrogen peroxide and the rate of NBT reduction is inhibited and can be measured spectrophotometrically at 560 nm. The rate of NBT reduction is calculated as percent inhibition, relative to the NBT reduction in the absence of the sample and the amount of protein causing 50% maximum inhibition is defined as one unit of activity. MnSOD activity was distinguished from CuZSOD activity using 5 mmol/L NaCN as described (6).

*Glutathione reductase (GR) activity*. GR activity was determined by the method or Ray and Prescott (7). Briefly, sample was diluted into a reaction mixture containing 70 mM KPi, pH 7.6, with 2.4 mM EDTA, 0.1% BSA, 94 □M

NADPH, and 1 mm GSSG. The loss of NADPH was assessed at room temperature by monitoring the change in absorbance at 340 nm.

*Thioredoxine reductase (TRR) activity*. TRR activity was determined using a commercially available kit (Sigma, CS0170), and was run as described.

*Glucose 6 phosphate dehydrogenase (G6PD)*. G6PD activity was measured using the method of Glock and McLean (8). Briefly, kidney homogenate was added to a solution containing 75 mM Tris, pH 8.0, with 0.75 mM MgCl2, 1.5 mM NADP+, and 1.2 mM glucose 6-phosphate. G6PD activity was determined by the rate of NADPH formation at 340 nm. To ensure the specificity of G6PD activity measurement, homogenates were additionally run in the presence of 1.2 mM 6-phosphogluconic acid, with and without glucose 6-phosphate.

*Citrate synthase activity*. Citrate synthase activity was measured using the protocol previously described (9). A sample volume equivalent to 50 micrograms of protein was combined with 0.5 mM oxaloacetic acid, and 0.1 mM dTNB. The formation of TNB was monitored at 412 nm every 30 minutes for 2 hours. Regression analysis was used to determine reaction rates.

*Mitochondrial ETC activities*. Measurements of the ETC complex activities were done as described previously (10). Complex I activity was assayed as the rate of rotenone-inhibitable NADH oxidation at 340 nm with 425 nm as reference. Samples were assayed with or without 200 µg/mL rotenone in working buffer containing 25 mM potassium phosphate buffer, 5 mM magnesium chloride, 2 mM potassium cyanide, 2.5 mg/mL BSA, 0.13 mM NADH, 200 µg/mL antimycin A, 7.5 mM coenzyme Q1. Complex II activity was assayed as the rate of reduction of 2,6-dichloroindophenol by coenzyme Q in the presence and absence of 0.2 M succinate at 600 nm. Samples were incubated with or without succinate in 25 mM potassium phosphate buffer, 5 mM magnesium chloride, 2 mM potassium cyanide, 2.5 mg/mL BSA for 10 minutes at 30 C. After incubation, 200 µg/mL antimycin A, 200 µg/mL rotenone, 5 mM DCIP, and 7.5 mM coenzyme Q1 were added to each cuvette and incubated for 1 min before reading absorbance rates. Complex III activity was assayed as the rate of cytochrome c reduction by coenzyme Q2 at 550 nm with 580 nm as reference. Samples were assayed in 25 mM potassium phosphate buffer, 5 mM magnesium chloride, 2 mM potassium cyanide, 2.5 mg/mL BSA, 0.5 mM n-dodecyl β-maltoside, 200 µg/mL rotenone, 1.5 mM cytochrome c, and 3.5 mM coenzyme Q2.

**Renal Tubule Epithelial Cell Isolation and Culture.** Primary Tubular Cells were obtained from 8-10-week-old MPC TubKO mice and WT controls. Mice were euthanized and kidneys were placed in cold-DPBS (Gibco, 14200-075). The capsule around the kidney was removed and the decapsulated kidney was finely minced with a sterile scalpel. Pooled renal cortices underwent digestion in HBSS containing 0.1% w/vol Type II Collagenase (Fisher Scientific, NC9693955), 1X non-essential amino acid (Gibco, 11-140-122), 0.2% BSA (Fisher Scientific, BP9706-100), and 0.5X penicillin/streptomycin (Gibco, 15-140-122) for 60 min at 37°C. The supernatant was collected and filtered using a sterile 70 μM Nylon Mesh cell strainer (Fisher Scientific, 22363548), and then the strained cell solution was centrifuged at 1200 RPM for 3 min. The cell pellet was resuspended in 12 ml of growth medium containing DMEM/Ham's F12 (1:1) (Gibco, 10567-014, Gibco 31765-035) supplemented with 100 U/mL penicillin, 100 mg/mL streptomycin, 1% fetal bovine serum (Gibco, 160000-0044), 1X non-essential amino acid solution, 5 mg/mL insulin, 5 mg/mL transferrin, 5 ng/mL selenite (Sigma Aldrich, I-1884), 36 ng/mL hydrocortisone (Sigma Aldrich, H0888), 4 pg/mL 3,3',5-Triiodo-L-thyronine (Sigma Aldrich, T0281), 10 ng/mL prostaglandin E1 (Sigma Aldrich, P8908), 15 ng/mL rhEGF (Sigma Aldrich, E9644), 0.5 ug/mL epinephrine (Sigma Aldrich, E4250) and 50 ug/mL L-ascorbic acid (Sigma Aldrich, A4544). The cell suspension was plated into T75 Flasks pre-coated with bovine collagen (Fisher Scientific, 12-565-349). The medium was changed every two days until cells reached 70 to 80% confluence after 6 to 7 days when cells were used experimentally or sub-cultured.

**Mitochondrial Isolation.** Freshly harvested and decapsulated kidney tissue was minced on ice and then homogenized in 10 ml of ice-cold buffer containing 70mM sucrose, 210mM d-mannitol, 1mM EGTA, 0.5% fatty acid free BSA, and 5mM HEPES pH 7.4. The homogenate was centrifuged at 800 x g for 10 minutes at 4°C. The supernatant was transferred to a new centrifuge tube and centrifuged at 8000 x g for 10 minutes at 4°C. The supernatant was discarded and the pellet, containing mitochondria, was gently resuspended in 10 ml of the homogenization buffer. The mitochondrial suspension was again centrifuged at 8000 x g. The supernatant was discarded, and the pellet was resuspended in 1 ml of the buffer above and centrifuged at 9000 x g for 10 minutes at 4°C. The supernatant was discarded, and the mitochondrial pellet was stored on ice until use.

**Metabolomic sample preparation.** Kidney tissue samples were lyophilized overnight, and metabolites were extracted in 18 volumes of a solvent solution:mg wet tissue weight. The solvent solution contained methanol/acetonitrile/water (2:2:1 V/V/V) with 1 ng/µl each of D4-Succinate, D8-Valine, D4-Citrate, 13C5-Glutamine, 13C5-Glutamate, 13C6-Lysine, 13C5-Methionine, 13C3-Serine, and 13C11-Tryptophan as internal standards (Cambridge Isotope). Tissues were homogenized using a ceramic bead mill homogenizer operated for 30 seconds at 6.45 MHz (Omni International). Homogenates were rotated at -20ºC for one hour and centrifuged at 20,000 x g at 4ºC for 10 minutes. The supernatant, referred to as metabolite extract, was collected for gas chromatography (GC)- and liquid chromatography (LC)- mass spectrometry (MS) analysis. A quality control (QC) sample was prepared by pooling equal volumes from each sample; the QC sample was aliquoted for GC- and LC-MS analysis as described below. The QC samples was analyzed at the beginning, end, and intermittently throughout GC- and LC-MS.

**GC-MS derivatization and analysis.** 150 µl of metabolite extract was transferred to an autosampler vial and dried to completeness using a Speedvac vacuum concentrator without heating (Thermo Fisher). Dried extracts were reconstituted in 24 µL of pyridine containing 11.4 mg/mL of methoxyamine (MOX). Samples were then vortexed for 10 min and then heated at 60°C for 90 minutes. Next, 16 µL of N-methyl-N-(trimethylsilyl) trifluoroacetamide (MSTFA) was added to the pyridine/MOX derivatized samples, pulse-vortexed, and heated at 60°C for 30 minutes. Analysis was conducted on a single quadrupole GC-MS (Trace 1300 GC + ISQ, Thermo Fisher). 1 µL of sample was injected into the GC (split ratio: 20-1; split flow: 24 mL/min, purge flow: 5 mL/min, Carrier mode: Constant Flow, Carrier flow rate: 1.2 mL/min). The oven method was as follows: 80°C for 3 minutes, ramp to 280°C at a rate of 20°C/min to a maximum temperature of 280°C and hold at 280°C for 8 minutes. For targeted metabolic profiling, the MS was operated from 3.90 to 21.00 minutes in SIM-mode using electron-ionization (-70eV).

**LC-MS analysis.** 400 µl of metabolite extract was transferred to a microcentrifuge and dried to completeness using a Speedvac vacuum concentrator without heating. Dried extracts were reconstituted in 40 µL acetonitrile/water (1:1 v/v), vortexed well, and centrifuged at 20,000 x g at 4ºC for 10 minutes. Finally, the cleared metabolite extracts were transferred to LC-MS autosampler vials for analysis. 2 µL of metabolite extracts were separated using a Millipore SeQuant ZIC-pHILIC (2.1 X 150 mm, 5 µm particle size) column with a ZIC-pHILIC guard column (20 x 2.1 mm) attached to a Thermo Vanquish Flex UHPLC. Mobile phase was comprised of Buffer A – 20 mM (NH4)2CO3, 0.1% NH4OH and Buffer B: acetonitrile. The chromatographic gradient was run at a flow rate of 0.150 mL/min as follows: 0–20 min—linear gradient from 80 to 20% Buffer B; 20–20.5 min—linear gradient from 20 to 80% Buffer B; and 20.5–28 min—hold at 80% Buffer B (11, 12). Data was acquired using a Thermo Q Exactive mass spectrometer operated in full-scan, polarity-switching mode with the spray voltage set to 3.0 kV, the heated capillary held at 275 °C, and the HESI probe held at 350 °C. The sheath gas flow was set to 40 units, the auxiliary gas flow was set to 15 units, and the sweep gas flow was set to 1 unit. MS data acquisition was performed in a range of m/z 70–1,000, with the resolution set at 70,000, the AGC target at 10e6, and the maximum injection time at 200 ms.

**REFERENCES**

1. Kim JY, Bai Y, Jayne LA, Abdulkader F, Gandhi M, Perreau T, et al. SOX9 promotes stress-responsive transcription of VGF nerve growth factor inducible gene in renal tubular epithelial cells. *J Biol Chem.* 2020;295(48):16328-41.
2. Kirita Y, Wu H, Uchimura K, Wilson PC, and Humphreys BD. Cell profiling of mouse acute kidney injury reveals conserved cellular responses to injury. *Proc Natl Acad Sci U S A.* 2020;117(27):15874-83.
3. Taylor CR, and Levenson RM. Quantification of immunohistochemistry--issues concerning methods, utility and semiquantitative assessment II. *Histopathology.* 2006;49(4):411-24.
4. Tietze F. Enzymic method for quantitative determination of nanogram amounts of total and oxidized glutathione: applications to mammalian blood and other tissues. *Anal Biochem.* 1969;27(3):502-22.
5. Griffith OW. Determination of glutathione and glutathione disulfide using glutathione reductase and 2-vinylpyridine. *Anal Biochem.* 1980;106(1):207-12.
6. Spitz DR, and Oberley LW. An assay for superoxide dismutase activity in mammalian tissue homogenates. *Anal Biochem.* 1989;179(1):8-18.
7. Ray LE, and Prescott JM. Isolation and some characteristics of glutathione reductase from rabbit erythrocytes (38548). *Proc Soc Exp Biol Med.* 1975;148(2):402-9.
8. Glock GE, and Mc LP. Further studies on the properties and assay of glucose 6-phosphate dehydrogenase and 6-phosphogluconate dehydrogenase of rat liver. *Biochem J.* 1953;55(3):400-8.
9. Barrientos A. In vivo and in organello assessment of OXPHOS activities. *Methods.* 2002;26(4):307-16.
10. Birch-Machin MA, Briggs HL, Saborido AA, Bindoff LA, and Turnbull DM. An evaluation of the measurement of the activities of complexes I-IV in the respiratory chain of human skeletal muscle mitochondria. *Biochem Med Metab Biol.* 1994;51(1):35-42.

11. Rauckhorst AJ, Borcherding N, Pape DJ, Kraus AS, Scerbo DA, and Taylor EB. Mouse tissue harvest-induced hypoxia rapidly alters the in vivo metabolome, between-genotype metabolite level differences, and (13)C-tracing enrichments. *Mol Metab.* 2022;66:101596.
12. Cantor JR, Abu-Remaileh M, Kanarek N, Freinkman E, Gao X, Louissaint A, Jr., et al. Physiologic Medium Rewires Cellular Metabolism and Reveals Uric Acid as an Endogenous Inhibitor of UMP Synthase. *Cell.* 2017;169(2):258-72 e17.
